# Macrophage-instructed GSDME couples glioblastoma cell-state plasticity with inflammatory cell death

**DOI:** 10.64898/2026.07.22.740157

**Authors:** Yihan Yang, Jonathan Sai-Hong Chui, Ellen Vervoort, Julie Van Nieuwenhuyze, Shikang Zhao, Carmen Bravo González-Blas, Kinga Anna Nowak, Sanket More, Kathryn A Jacobs, Aaron Ziani Zeryouh, Nikolina Dubroja, Raf Sciot, An Coosemans, Massimiliano Mazzone, Fredrik De Smet, Steven De Vleeschouwer, Patrizia Agostinis

**Affiliations:** Cell Death Research & Therapy, VIB Center for Cancer Biology, Leuven, Belgium; Department of Cellular and Molecular Medicine, KU Leuven, Leuven, Belgium; Research Group Experimental Neurosurgery and Neuroanatomy, Department of Neurosciences, KU Leuven, Leuven, Belgium; Leuven Institute for Single-Cell Omics (LISCO), Leuven, Belgium; Laboratory for Precision Cancer Medicine, Translational Cell and Tissue Research Unit, Department of Imaging & Pathology, KU Leuven, Leuven, Belgium; Laboratory of Tumor Immunology and Immunotherapy, Department of Oncology, Leuven Cancer Institute, KU Leuven, Belgium; Laboratory of Tumor Inflammation and Angiogenesis, VIB-KU Leuven Center for Cancer Biology, Leuven, Belgium; Department of Oncology, KU Leuven, Leuven, Belgium; Department of Pathology, KU Leuven and University Hospitals Leuven, Leuven, Belgium; Department of Neurosurgery, University Hospitals Leuven, Leuven, Belgium; Leuven Cancer Institute (LKI), KU Leuven, Leuven, Belgium; Leuven Brain Institute (LBI), KU Leuven, Leuven, Belgium

**Author notes:** Correspondence (S.D.V), (P.A.).

**Keywords:** Glioblastoma, gasdermin E, tumor-associated macrophages, tumor cell plasticity, macrophage-tumor cell crosstalk, spatial tumor microenvironment, mesenchymal cell state, inflammatory cell death

## Abstract

Glioblastoma (GBM) cell states reflect spatial microenvironmental interactions. Here, using COMET spatial proteomics and RNAscope across multiregional human GBM tissue, spanning tumor cores with pseudopalisading regions, infiltrative margins and peripheral regions, together with multiplex spatial profiling of 202 specimens from 33 patients with matched primary and recurrent tumors, we identify full-length gasdermin E (GSDME-FL) as a macrophage-instructed regulator of malignant cell plasticity. Integration with single-cell transcriptomics and functional perturbation shows that GSDME-high tumor cells localize to macrophage-rich perivascular niches and are associated with delayed recurrence and longer patient survival. Mechanistically, macrophage-derived S100A4 activates EGFR-Sp1 signaling to induce GSDME-FL in neighboring GBM cells. GSDME-FL restrains hypoxia-associated mesenchymal plasticity and shapes macrophage-induced tumor state transitions independently of caspase activation. During immunogenic cell death (ICD), cleaved GSDME promotes pre-lytic swelling, early ATP efflux, and the release of canonical ICD-associated cytokines and chemokines. NLRP3 signaling further supports ATP release and licenses macrophage phagocytosis of dying GBM cells, whereas NINJ1-dependent membrane rupture enables terminal HMGB1 release. Vaccination with ICD-treated glioma cells elicits tumor rejection in a prophylactic intracranial challenge model. Notably, co-expression of GSDME and NINJ1 in macrophage-rich perivascular niches provides a spatial correlate of pathway convergence in patient tumors. Thus, our findings define how a spatial macrophage niche induces GSDME-FL, thereby coupling malignant cell state plasticity to the inflammatory properties of GBM cell death.

## Introduction

Glioblastoma (GBM) is the most aggressive primary brain tumor in adults and is characterized by diffuse infiltration, rapid progression, limited treatment responses and almost inevitable recurrence. Standard treatment, comprising surgery followed by radiotherapy and temozolomide (TMZ), provides only a modest survival benefit, with a median survival of approximately 15 months^1^. The dismal prognosis reflects the remarkable heterogeneity and plasticity of GBM^2^, which enable it to rapidly adapt to metabolic and therapeutic stress. Single-cell transcriptomic and lineage-tracing studies have established that GBM is a dynamic ecosystem of interconnected malignant-cell states rather than a collection of discrete molecular subtypes^2^. Individual GBM cells can occupy and transition between, at least four major programs: oligodendrocyte-progenitor-cell-like (OPC-like), neuronal-progenitor-cell-like (NPC-like), astrocyte-like (AC-like) and mesenchymal-like (MES-like). A recent study expanded the repertoire of malignant programs, supporting a multidimensional, context-dependent network of interconnected cell states^3^.

This plasticity is further compounded by the spatial architecture of GBM and the dynamic interactions between cancer cells and the tumor microenvironment (TME), fostering a profoundly immunosuppressive milieu characterized by poor cytotoxic T cell infiltration. In line with this, GBM develops within a myeloid-rich microenvironment in which microglia and macrophages constitute a large fraction of tumor cellularity and regulate invasion, angiogenesis, immune suppression and therapy response^2,4–6^. However, resident microglia and monocyte-derived macrophages differ in ontogeny, spatial localization (invasive front versus hypoxic/perivascular cores) and functional programs^7,8^. These distinctions highlight the need to resolve GBM-myeloid interactions as spatially and functionally specialized niches. How signals from these niches are encoded within malignant cells remains incompletely understood.

The inflammatory microenvironment is increasingly recognized as a central modulator of GBM biology, cancer stem cell behavior and therapy-induced cell death. These processes can both amplify antitumor immunity or promote tumor growth, depending on the context^9–13^. However, the molecular executors linking inflammation to cell death susceptibility and tumor adaptation in GBM are poorly defined. Members of the gasdermin (GSDM) family are key effectors of pyroptosis, an inflammatory form of regulated cell death^14–19^. Upon activation by caspases, the N-terminal domain of GSDMs oligomerizes in the plasma membrane, forming 20-40nm pores that promote ionic imbalance and swelling, while facilitating the efflux of pro-inflammatory cytokines and ultimately lytic cell death^17,20,21^, bridging innate sensing to immune activation and tissue damage. Among the GSDM family members, inflammasome-mediated cleavage of GSDMD represents the best characterized pore-forming executor of pyroptosis, whereas GSDME cleavage downstream of caspase-3 activation contributes to convert apoptosis into pyroptosis-like cell death^17–19^. Accumulating evidence nevertheless indicates that proteolytic cleavage does not fully account for GSDM biology and that full-length GSDMs can exert additional cellular functions^20,22^. Specific GSDMs, including GSDMB, have been shown to participate in tissue repair and epithelial regeneration, independent of pyroptosis ^23^. These noncanonical activities raise the possibility that full-length GSDMs may influence cellular signaling and stress adaptations in a tumor context-dependent way.

Although GSDMs have been implicated in host defense, inflammation, and tissue homeostasis^24^, a recent study has suggested a pro-tumorigenic role of GSDM expression in glioma^25^. Given the inflammatory, hypoxic and metabolically stressed nature of the GBM microenvironment, GSDMs are well-positioned to integrate cell death, inflammatory signaling and tumor adaptation in this setting. However, how GSDME expression is regulated within spatially defined GBM niches and how this regulation influences malignant-cell plasticity are unknown. It is also unknown whether a niche-imprinted GSDME state in viable tumor cells subsequently shapes inflammatory consequences during cell death.

Here, by integrating spatial profiling of primary and recurrent human GBM with single-cell transcriptomics and functional perturbations, we identify a macrophage-associated perivascular niche in which tumor cells express predominantly full-length GSDME. GBM cells promote a scavenger-like macrophage state that, in turn, induces tumor-cell GSDME through S100A4-dependent EGFR-Sp1 signaling, establishing reciprocal GBM-macrophage crosstalk. Under non-lytic conditions, loss of GSDME biases tumor cells toward hypoxia-associated mesenchymal programs and alters their response to macrophage-derived cues. In a therapeutic stress context, GSDME, NLRP3 signaling and NINJ1-associated membrane rupture coordinate complementary inflammatory outputs from dying GBM cells. Together, these findings show how macrophage-derived signals sustain a predominantly full-length GSDME pool that modulates tumor cell plasticity, whereas proteolytic activation enables GSDME to shape inflammatory signaling during cell death.

## Results

### Integrative spatial analysis links GSDME expression in GBM cells to macrophage proximity

GSDME has recently been reported to be highly expressed in GBM, but the mechanisms sustaining its expression in malignant cells remain undefined^25^. To delineate the mechanisms governing GSDME expression in GBM, we analyzed TCGA pan-cancer datasets and examined immunohistochemistry data from the Human Protein Atlas (HPA). These analyses confirmed that GSDME is elevated in GBM relative to most other solid tumors **(Supplementary Figure 1A, left)**. Within the gasdermin family, GSDME showed the strongest GBM-associated enrichment relative to normal brain tissue, followed by a more modest enrichment of GSDMD **(Supplementary Figure 1A, right)**. At the protein level, GSDME was consistently detected in patient-derived cell lines (PDCLs), including the predominantly classical primary and recurrent cell lines CME037 and CME038 and a primarily mesenchymal primary cell line (CME014) **(Supplementary Figure 1B and C)**. In contrast, GSDMD expression was largely restricted to the CME014 **(Supplementary Figure 1C)**. Analysis of HPA cohorts further showed that GSDME expression was predominantly detected in high -grade gliomas **(Supplementary Figure 1D)**. These observations prompted us to focus on the mechanisms and functional consequences of GSDME upregulation in GBM, while not excluding a potential contributory role of GSDMD.

Single cell and spatial transcriptomic analyses have established that GBM exhibits pronounced cellular and transcriptional heterogeneity, underpinned by transcriptional programs representing distinct tumor cell states^2,26^. Given this context-dependent plasticity and the strongly non-cell-autonomous nature of GBM, we hypothesized that GSDME expression may be influenced not only by tumor-intrinsic programs but also by spatial interactions within the tumor microenvironment (TME).

To test this, we integrated high-plex COMET spatial proteomics with RNAscope in situ hybridization to resolve GSDME-expressing malignant cells and their neighboring microenvironment at single cell resolution. We applied COMET to 16 GBM tissue sections representing multiple tumor regions, using a marker panel that captured malignant cells and major non-malignant cell types **(Figure 1A, Supplementary Figure 1E, Supplementary Figure 2 and 3)**. These sections were obtained from the tumors from which CME037, CME038 and CME014 PDCLs were established. In this cohort, GSDME expression was largely restricted to tumor tissue rather than adjacent normal tissue **(Supplementary Figure 1F)**. At the cellular level, although GSDME was detectable in macrophages, in line with prior reports in other contexts^27^, GBM cells exhibited robust and concordant GSDME expression at both the RNA (RNAscope) and protein (COMET) levels **(Supplementary Figure 1G)**. We therefore focused subsequent analyses on tumor cell-intrinsic GSDME.

**Figure 1.**
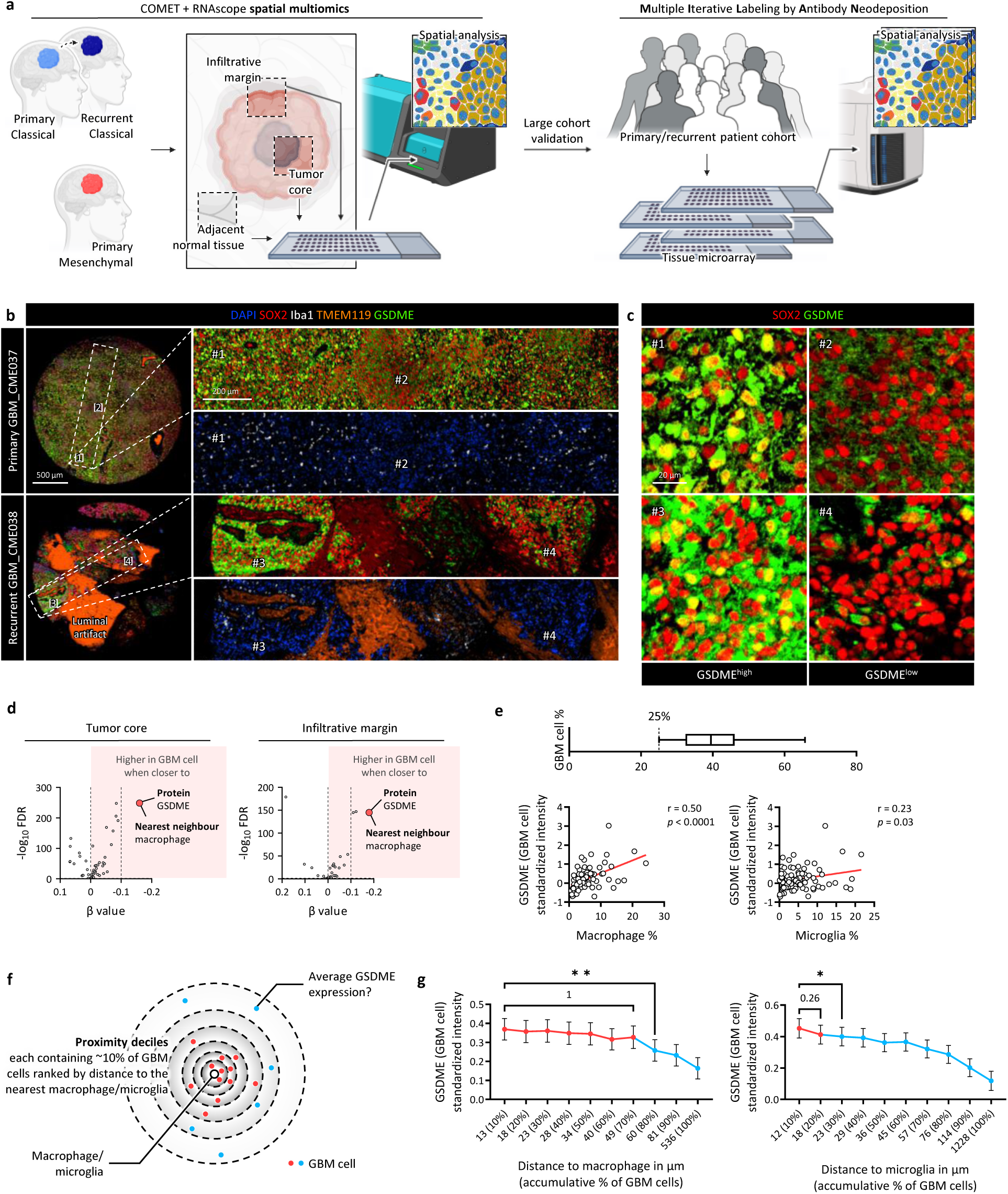
Macrophage proximity is associated with elevated GSDME expression in GBM cells. **(A)** Schematic of sample cohorts and spatial omics approaches. **(B)** Multiplexed protein images of SOX2, Iba1, TMEM119, and GSDME in matched primary (CME037) and recurrent (CME038) GBM bulk tissues profiled using the COMET platform. **(C)** Magnified views of ROIs 1-4 indicated in **(B)**, showing SOX2 and GSDME staining in GSDME-high (ROIs 1 and 3) and GSDME-low (ROIs 2 and 4) regions. **(D)** Volcano plots showing associations between GBM-cell protein expression and proximity to neighboring non-malignant cell types in the tumor core (left) and infiltrative margin (right). Each dot represents one protein-neighboring cell type pair. Negative distance coefficients, plotted to the right, indicate higher expression in GBM cells proximal to the indicated neighboring cell type. Positive distance coefficients, plotted to the left, indicate higher expression in distal GBM cells. Significance is shown as -log_10_(FDR). **(E)** Pearson correlation between macrophage or microglia abundance and standardized GSDME expression per GBM cell per image. Images in which GBM cells accounted for ≥25% of segmented cells were included. **(F)** Schematic of the proximity analysis. Macrophages or microglia were used as index cells, and surrounding GBM cells were ranked by distance to the nearest index cell and partitioned into deciles, with each decile containing approximately 10% of GBM cells. GSDME expression per GBM cell was then compared across deciles. **(G)** GSDME expression per GBM cell across distance deciles centered on macrophages (left) or microglia (right). Statistical significance is indicated as *p < 0.05, **p < 0.01.

High-plex COMET spatial proteomics revealed that GSDME-expressing tumor cells were mixed with macrophages across the sampled regions of paired primary and recurrent tumors corresponding to the predominantly classical PDCLs **(Figure 1B and C)**. Spatial neighbor analysis showed that GSDME was selectively elevated in GBM cells located near macrophages within the tumor core and infiltrative margin **(Figure 1D)**. A mesenchymal GBM specimen corresponding to CME014 presented a similar organization, with high GSDME expression in tumor areas containing abundant macrophages **(Supplementary Figure 1H)**. GSDME^high^ malignant cells localized to macrophage-enriched niches, whereas resident microglia were largely absent from these regions **(Figure 1B and Supplementary Figure 1H)**. Hence, a contribution from microglia could not be excluded yet, owing to their low abundance in this cohort.

Because GSDME can be activated by caspase-3-mediated cleavage^19^, we next asked whether GSDME-positive regions reflected areas associated with active cell death. Although pro-caspase-3 was detectable, cleaved caspase-3-positive cells were infrequent and rarely detected in GBM cells by quantitative analysis **(Supplementary Figure 1I)**. Together, this spatial analysis indicated that malignant cells found in a macrophage-rich context express full-length GSDME without evidence of local caspase-3 activation.

To validate the GBM-macrophage association in a larger cohort with substantial microglial representation **(Supplementary Figure 1M)**, while enabling assessment of the contribution of microglia to GSDME expression in GBM cells, we applied multiple iterative labeling by antibody neodeposition (MILAN)-based spatial proteomics^28^ to a total of 202 GBM specimens from 33 patients with matched primary and recurrent disease **(Figure 1A and Supplementary Figure 1J-M)**. We examined the association between GSDME expression in GBM cells and myeloid cell abundance across this cohort. GSDME intensity in malignant cells correlated with tumor-associated macrophage (TAM) abundance but showed little association with microglia **(Figure 1E)**. To resolve this relationship spatially, we quantified tumor-myeloid cell proximity using nearest-neighbor analysis^29^. For each tumor section, the Euclidean distance from every malignant cell to the nearest macrophage or microglial cell was computed. Malignant cells were then stratified into proximity deciles, and mean GSDME intensity was calculated for each decile at the image level to generate a distance-resolved spatial gradient **(Figure 1F)**. Across tumor sections, GSDME intensity in malignant cells was maintained within ∼50 μm of macrophages and declined with increasing distance **(Figure 1G, left)**. In contrast, GSDME intensity decreased within ∼18 μm of microglia, plateaued at intermediate distances, and showed a further reduction beyond ∼45 μm **(Figure 1G, right)**, suggesting that the association between GSDME^high^ malignant cells and macrophages extends more broadly across the tumor parenchyma than the association with microglia.

Together, these data reveal that both primary and recurrent GBM harbor GSDME^high^ tumor cells are predominantly positioned near infiltrating macrophages rather than resident microglia.

### Macrophage-derived S100A4 drives GSDME expression via an EGFR-Sp1 axis

Our spatial mapping showing that GSDME^high^ tumor cells preferentially localized near macrophages within intratumoral niches suggested a functional interaction. To examine whether macrophage proximity could directly regulate GSDME expression in tumor cells, we reanalyzed published single cell RNA sequencing (scRNA-seq) data from PDCLs cultured alone or in the presence of monocyte-derived macrophages from healthy donors^30^. This analysis revealed a consistent increase in *GSDME* transcript abundance following co-culture, whereas *GSDMD* was not markedly induced **(Supplementary Figure 4A-C)**.

To validate this effect in a tractable model, we cultured PDCLs, obtained from the same matched primary and recurrent GBM samples used in our spatial analysis **(Figure 1B-D)**, either alone or with PMA-differentiated THP-1-derived macrophage-like cells (THP-1-macrophages; **Figure 2A**). Quantitative PCR (qPCR) and immunoblotting confirmed increased GSDME expression at both the mRNA and protein levels across PDCLs **(Figure 2B,C and Supplementary Figure 4D)** with different GBM subtypes **(Supplementary Figure 1B)**. In contrast, no GSDMD upregulation was observed under the same conditions **(Figure 2B and Supplementary Figure 4A-D)**, and monocytic THP-1 cells did not increase GSDME expression in GBM cells either **(Supplementary Figure 4E)**. Co-culture with the microglial cell line HMC3 also failed to induce GSDME upregulation, a result corroborated by external bulk RNA-seq data of GBM cells co-cultured with human microglia^31^ **(Supplementary Figure. 4F and 4G)**. While not excluding a potential function of microglia, this finding argues against their major contribution to GSDME upregulation, consistent with our spatial proteomics analysis **(Figure 1E and 1G)**.

**Figure 2.**
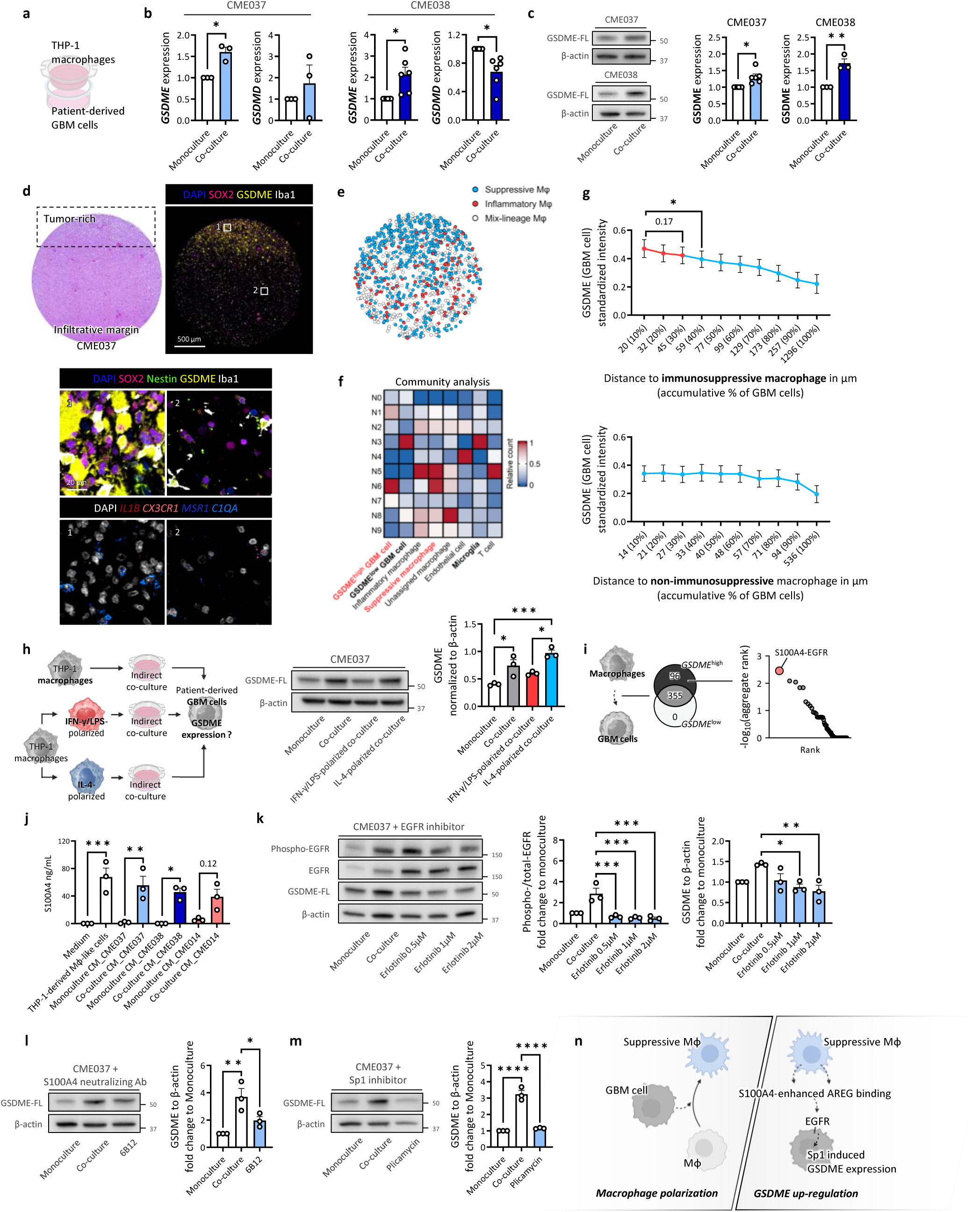
Immunosuppressive macrophages promote GSDME expression through S100A4-EGFR-Sp1 signaling. **(A)** Schematic of the indirect co-culture system. **(B)** qPCR analysis of *GSDME* and *GSDMD* expression in PDCL CME037 and CME038 cultured alone or indirectly co-cultured with THP-1-derived macrophage-like cells. **(C)** Immunoblots of GSDME expression in GBM PDCLs after indirect co-culture with THP-1-derived macrophage-like cells. **(D)** COMET images of tumor tissue corresponding to PDCL CME037, showing SOX2, Nestin, Iba1, GSDME, and DAPI, together with RNA staining for *IL1B*, *CXCR1*, *C1QA*, and *MSR1*. **(E)** Scatter plot showing the spatial distribution of annotated macrophage subtypes in the tissue shown in **(D)**. **(F)** Heatmap showing the relative count of each cell type across the niches identified by community analysis. **(G)** GSDME expression per GBM cell across distance deciles using immunosuppressive macrophages (left) or non-immunosuppressive macrophages (right) as index cells. **(H)** Schematic of the indirect co-culture system using polarized macrophages and GBM PDCLs (left). Representative immunoblot of GSDME expression under the indicated culture conditions (middle) and quantification of normalized GSDME expression (right). **(I)** Cell-cell communication analysis identifying macrophage-to-GBM PDCL ligand-receptor pairs conserved across all PDCLs. Specific ligand-receptor pairs from macrophages to *GSDME*^high^ GBM cells are ranked by -log_10_(aggregated rank). **(J)** ELISA quantification of S100A4 in macrophage-like cell monoculture, GBM PDCL monoculture, and indirect co-culture conditions. **(K)** Immunoblots and quantification of GSDME expression following S100A4 neutralizing antibody treatment in the indirect co-culture system. **(L)** Immunoblots and quantification of the phosphorylated-to-total EGFR ratio and GSDME expression after erlotinib treatment in the indirect co-culture system. **(M)** Immunoblots and quantification of GSDME expression after plicamycin treatment in the indirect co-culture system. **(N)** Schematic model showing that GBM cells polarize tumor-associated monocyte-derived macrophages toward an immunosuppressive state, which sustains GSDME expression in tumor cells through S100A4 and amphiregulin secretion and EGFR-SP1 signaling. Data are shown as mean ± SEM. Each dot in the bar plots represents an individual replicate. Statistical significance is indicated as *p < 0.05, **p < 0.01, ***p < 0.001.

In our co-culture systems **(Figure 2A-C and Supplementary Figure 4A-D)**, GSDME upregulation required prolonged macrophage exposure (approximately 4 days), suggesting that sustained macrophage-tumor cell interaction, possibly involving macrophage phenotypic reprogramming, rather than acute signaling alone, drove this effect. Consistent with this, prior scRNA-seq analysis showed progressive polarization of macrophages toward immunosuppressive states during GBM-macrophage co-culture^30^. Applying a recent myeloid cell-state classification^32^, we observed a consistent enrichment of the scavenger suppressive phenotype in macrophages co-cultured with PDCLs, whereas complement immunosuppressive and systemic inflammatory programs varied among cell lines **(Supplementary Figure 4H)**.

RNAscope analysis corroborated these findings, showing that macrophages adjacent to GSDME^high^ GBM cells co-expressed high levels of *MSR1* (*CD204*), a marker of scavenger-suppressive macrophages, and *C1QA*, a marker of complement-suppressive macrophages, whereas *IL1B* and *CX3CR1*, characteristic of systemic inflammatory macrophages and inflammatory microglia, were largely absent **(Figure 2D)**. Spatial reconstruction of macrophage phenotypes confirmed this pattern **(Figure 2E)**.

Unbiased community analysis of tissues from the COMET cohort further supported these findings. Among the identified spatial communities, niche 6 (N6) was characterized by the co-localization of GSDME^high^ GBM cells with immunosuppressive macrophages expressing canonical suppressive markers, including *CD163, MSR1* and *C1QA*. In contrast, niche 3 (N3) comprised predominantly GSDME^low^ GBM cells co-localized with microglia **(Figure 2F)**. In the larger MILAN cohort **(Supplementary Figure 1M and Supplementary Figure 4I)**, GSDME^high^ tumor cells were preferentially located in proximity, within ∼45 μm, to immunosuppressive macrophages **(Figure 2G)**. By contrast, regions adjacent to non-immunosuppressive macrophages showed lower GSDME expression and no comparable spatial association **(Figure 2G)**. Notably, GSDME upregulation following co-incubation with THP-1-macrophages was recapitulated by IL-4-polarized cells, but not by THP-1-macrophages induced toward an IFNγ/LPS-polarized phenotype under the same conditions **(Figure 2H)**.

We next hypothesized that GBM-driven macrophage reprogramming, in turn, promotes GSDME expression in tumor cells via soluble factors, establishing a reciprocal reinforcement loop. To identify potential mediators, we applied LIANA, a computational framework that integrates multiple cell-cell communication inference methods with consensus scoring across ligand-receptor databases^33^. LIANA identified 96 macrophage-to-GBM signaling interactions enriched in GSDME^high^ GBM cells, with S100A4-EGFR ranking highest **(Figure 2I)**. The calcium-binding protein S100A4 binds multiple EGFR-family ligands, with preferential interaction with amphiregulin (AREG), and potentiates EGFR activation^34^. In our spatial multiomics cohort, S100A4 RNA and protein were most expressed in macrophages and T cells **(Supplementary Figure 4J)**, consistent with human glioma scRNA-seq data^35^. Because T cells are scarce in GBM, macrophages likely account for the dominant source of S100A4, supporting our focus on macrophage-derived secretion. Accordingly, both S100A4 and AREG were detected in the macrophage-GBM co-culture supernatants, whereas they were absent in GBM monocultures **(Figure 2J and Supplementary Figure 4K)**.

To test the causal role of macrophage-derived S100A4 in GSDME upregulation, we performed co-culture in the presence of an S100A4-neutralizing antibody. Neutralization of S100A4 using the 6B12 reduced THP-1-macrophage-induced GSDME upregulation in PDCLs **(Figure 2L)**.

We next asked whether this response required EGFR activity. The PDCLs used for mechanistic validation did not harbor EGFRvIII, a constitutively active EGFR mutant, allowing us to interrogate ligand-responsive EGFR signaling **(Supplementary Figure 4L)**. Non-cytotoxic doses of the EGFR inhibitor erlotinib **(Supplementary Figure 4M)** reduced both EGFR phosphorylation and GSDME upregulation induced by the co-culture with THP-1-macrophages **(Figure 2K)**. By contrast, inhibiting tonic EGFR-STAT3 signaling in GBM monocultures did not reduce GSDME levels **(Supplementary Figure 4N)**, suggesting that STAT3 is not a primary regulator of GSDME expression. Consistently, pharmacological inhibition of JAK-STAT signaling also failed to attenuate macrophage-induced GSDME upregulation in PDCL co-cultures **(Supplementary Figure 4O)**, suggesting that macrophages engage alternative signaling downstream of the S100A4-EGFR axis. GSDME expression in other cancers is regulated by the transcription factor Sp1^36^, which is directly regulated by ERK downstream of EGFR^37^. Consistent with convergence on this signaling axis, plicamycin attenuated macrophage-induced GSDME expression in the adherent cell fraction tested by immunoblotting **(Figure 2M)**.

Altogether, these findings suggest the existence of a feed-forward interaction in which GBM cells promote a scavenger-like, immunosuppressive macrophage state that, in turn, drives GSDME upregulation in tumor cells via a S100A4-EGFR-Sp1 axis **(Figure 2N)**, within spatially restricted GBM-macrophage areas.

### GSDME^high^ tumor cells delineate spatially organized GBM architectures

We next asked whether GSDME^high^ tumor cells represent a dispersed molecular state or are preferentially embedded within organized tissue structures in vivo, and how they map onto the spatial architecture of patient GBM. Therefore, we integrated GSDME spatial profiles with neuropathologist-defined histopathological annotations across the tumor core and infiltrative margin. These annotations captured major GBM tissue states, including pseudopalisading necrotic areas, a defining hallmark of GBM, as well as vascular-rich regions, tumor cell-dense hypoxic regions, and hemorrhagic/mixed regions.

Within these annotated regions, GSDME^high^ GBM cells were frequently positioned near macrophages and vascular structures lined by endothelial cells **(Figure 3A)**, many of which contained erythrocytes on matched H&E sections and showed corresponding luminal artifacts in COMET images^38^. In line with this spatial pattern, perivascular macrophages expressed the suppressive macrophage markers *HMOX1*, *MSR1* and *C1QA*, as well as S100A4 at both the RNA and protein levels **(Figure 3B)**. This organization was observed across disease stages, suggesting that GSDME upregulation is maintained within actively expanding tumor regions rather than being restricted to a transient local state **(Figure 3C)**.

**Figure 3.**
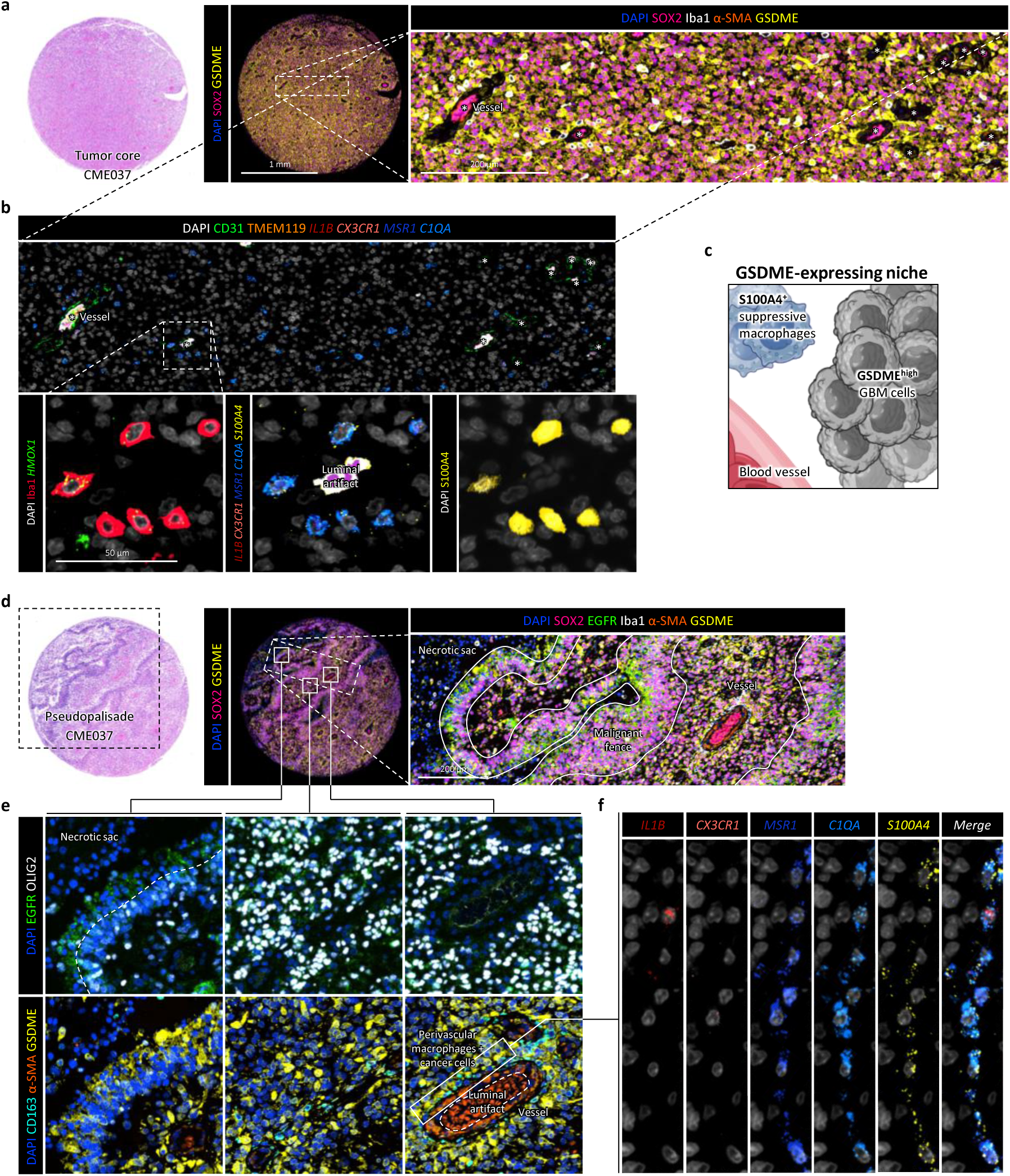
GSDME^high^ tumor cells localize to vascular-S100A4^+^ suppressive macrophage niches in GBM. **(A)** H&E image of a tumor core region from tumor tissue corresponding to PDCL CME037 (left) and corresponding COMET multiplexed staining for SOX2, GSDME, and DAPI, with a magnified region of interest showing SOX2, Iba1, α-SMA, GSDME, and DAPI. **(B)** Region of interest from **(A)** showing CD31, TMEM119, *IL1B*, *CX3CR1*, *MSR1*, *C1QA*, and DAPI (top), and a magnified image showing perivascular Iba1^+^ cells together with *HMOX1*, *IL1B*, *CX3CR1*, *MSR1*, *C1QA*, *S100A4* RNA, and S100A4 protein (bottom). **(C)** Schematic of the GSDME-expressing niche, comprising S100A4+ immunosuppressive macrophages, GSDME^high^ GBM cells, and blood vessels. **(D)** H&E image of a pseudopalisading region from tumor tissue corresponding to PDCL CME037 (left) and corresponding COMET multiplexed staining for SOX2, EGFR, Iba1, α-SMA, and GSDME in the same tissue region. **(E)** Region of interest from **(B)** showing EGFR and OLIG2 staining (top) and CD163, α-SMA, and GSDME staining (bottom). **(F)** Magnified region of interest from **c** showing RNAscope staining for *IL1B*, *CX3CR1*, *MSR1*, *C1QA*, and *S100A4*.

Because pseudopalisading necrosis represents a characteristic GBM architecture, we next examined how GSDME relates to this structure. A published analysis of the Ivy Glioblastoma Atlas Project (GAP) in the context of pyroptosis suggested that *GSDME* RNA expression is elevated in pseudopalisading regions^25^. However, single cell-resolved spatial profiling revealed a more nuanced organization. The hypercellular pseudopalisading rim itself, composed of densely packed tumor cells, showed low GSDME protein expression and contained relatively few adjacent macrophages **(Figure 3D and 3E)**. Instead, the elevated regional GSDME signal was attributable to tumor cells located outside the rim, in proximity to neighboring blood vessels surrounded by macrophages and multilayered GSDME^high^ GBM cells **(Figure 3D and 3E)**. RNAscope further showed that macrophages within this GSDME^high^ niche expressed *MSR1* and *C1QA*, consistent with an immunosuppressive phenotype, and were strongly positive for *S100A4*, supporting their potential role in sustaining GSDME expression in adjacent tumor cells **(Figure 3F)**.

Because retrievable far-invasive regions were limited in our COMET cohort, we complemented our analysis with two independent public scRNA-seq datasets containing this compartment. Consistent with our spatial data, tumor cell GSDME expression was significantly higher in the tumor core **(Supplementary Figure 5A, 5D and 5E)**, where immunosuppressive macrophages were more abundant and expressed high levels of *S100A4* and *AREG* **(Supplementary Figure. 5B and 5F-H)**. In contrast, the invasive front was enriched for inflammatory microglia and showed lower *GSDME* expression in tumor cells **(Supplementary Figure. 5A and 5E)**.

Together, the integration of spatial and scRNA-seq analyses indicates that GSDME expression in GBM cells is organized within recurrent vascular-macrophage-associated tissue structures in human tumors. This raised the question of whether GSDME simply reflects microenvironmental heterogeneity or functionally contributes to the GBM cell states that emerge within these architectures.

### Full-length GSDME coordinates mesenchymal transitions and stem-like states

To address the question experimentally, we used PDCLs derived from matched primary and recurrent GBM samples used in our previous functional assays and spatial analysis. Despite a degree of intra-model heterogeneity, each PDCL exhibited a dominant transcriptional phenotype **(Supplementary Figure 1B)**. We then generated CRISPR-Cas9-mediated GSDME knockout (KO) in these models **(Figure 4A)**, performed scRNA-seq, and applied gene set enrichment analysis (GSEA) using a curated pan-cancer metaprogram (MP) framework incorporating glioma-specific transcriptional programs^39^ **(Figure 4B and Supplementary Figure 6A)**.

**Figure 4.**
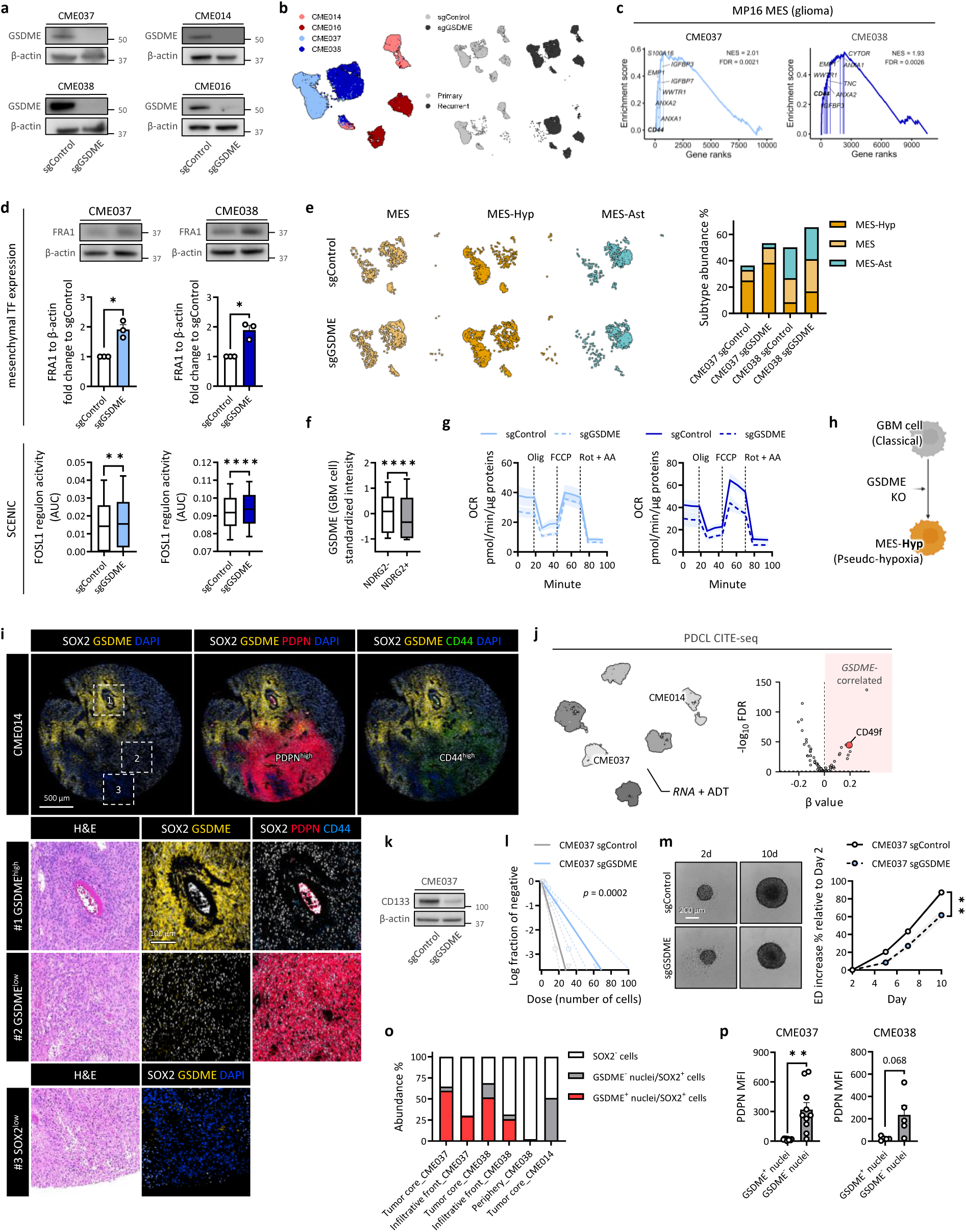
GSDME buffers hypoxic mesenchymal phenotype, enforcing stemness trade-off. **(A)** Immunoblots validating CRISPR-Cas9-mediated GSDME knockout (KO) in PDCL models. **(B)** scRNA-seq analysis of sgControl and sgGSDME PDCLs visualized on a UMAP embedding, showing PDCL origin (left), genotype (right top), and GBM stage (right bottom). **(C)** GSEA showing upregulation of the malignant meta-program MP16, MES (glioma), in sgGSDME CME037 and CME038. Labeled genes represent selected biologically relevant leading edge genes. **(D)** Immunoblot and quantification of FRA1, encoded by FOSL1, in sgControl and sgGSDME cells (top). FOSL1 regulon activity scores comparing sgControl and sgGSDME cells are shown below. **(E)** Mesenchymal subtypes defined by Greenwald et al. projected onto the UMAP embedding (left), distribution of these subtypes in sgControl and sgGSDME cells (right). **(F)** Standardized GSDME expression per GBM cell comparing normoxic and hypoxic GBM cells from the COMET™ staining. **(G)** Seahorse analysis comparing oxygen consumption rate (OCR) in sgControl and sgGSDME PDCLs. **(H)** Schematic showing transition of AC-like/classical GBM cells to mesenchymal, (pseudo-)hypoxic-prone GBM cells after GSDME KO. **(I)** COMET image of tumor tissue corresponding to the mesenchymal-dominant PDCL CME014, showing SOX2, GSDME, PDPN, CD44, and DAPI, with magnified views of the indicated ROIs. **(J)** CITE-seq analysis of PDCLs under basal conditions visualized on a UMAP embedding (left) and volcano plot (right) showing effect sizes (β) from single-cell linear mixed-effects models for each antibody-derived tag (ADT) comparing *GSDME*^hi^ versus *GSDME*^lo^ cells. Significance is shown as -log_10_(FDR). **(K)** Immunoblot of CD133 in sgControl and sgGSDME CME037. **(L)** Limiting dilution assay GBM PDCL cluster-forming frequency between sgControl and sgGSDME conditions. **(M)** Gliomasphere assay comparing sgControl and sgGSDME PDCL, with quantification of sphere morphology shown as percentage-change kinetics in equivalent diameter. **(N)** Bar plots showing the composition of GBM cells classified according to nuclear GSDME positivity. **(O)** PDPN mean fluorescence intensity (MFI) in GBM cells stratified by nuclear GSDME positivity. Data are shown as mean ± SEM. Each dot in the bar plots represents an individual replicate. Statistical significance is indicated as *p < 0.05, **p < 0.01, ***p < 0.001.

Across PDCLs, GSDME loss reprogrammed tumor cell states toward mesenchymal-associated programs, most prominently in models that were classical-dominant in the GSDME-intact control condition **(Figure 4C, Supplementary Figure 6A and 6B)**. In the mesenchymal-dominant models, the same trend was observed but was less pronounced **(Supplementary Figure 6A and 6B)**. Our previous work and that of others identified FOSL1 as a key transcriptional regulator of the mesenchymal phenotypic transition in GBM^40–42^. In this context, GSDME KO was accompanied by increased expression of FRA1, encoded by *FOSL1*, together with elevated regulon activity by SCENIC analysis **(Figure 4D)**.

We therefore focused on the classical-dominant models to resolve the mesenchymal programs induced by GSDME loss. To do this, we applied a spatially informed GBM classification that separates mesenchymal states into three partially overlapping metaprograms: a canonical MES-like program, a hypoxia-associated MES-Hyp program, and an astrocytic-like MES-Ast program^26^. Using this framework, we found that GSDME loss induced a shift toward MES-like and MES-Hyp modules, whereas the MES-Ast program was not substantially affected **(Figure 4E)**. These data suggest that GSDME loss promotes a hypoxia-biased mesenchymal transition rather than a broad activation of all mesenchymal states. GSDME-deficient cells showed increased activity of the hypoxia metaprogram (MP) **(Supplementary Figure 6A)** and upregulation of hypoxia-associated genes **(Supplementary Figure 6C)**. In line with these transcriptional changes, our spatial COMET analysis showed significantly reduced GSDME protein expression in *NDRG2*^+^ GBM cells in vivo **(Figure 4F)**. Functionally, GSDME-deficient cells exhibited reduced mitochondrial respiration, consistent with metabolic reprogramming toward a hypoxia-like state **(Figure 4G)**. Interestingly, GSDME loss also increased interferon- and MHC-II-associated programs, including interferon-stimulated genes across all models **(Supplementary Figure 6D-F)**. Of note, in several cancer types, mesenchymal state is linked to inflammatory and antigen-presentation-like programs and may promote immune evasion through STAT1-modulated immune-checkpoint regulation^43,44^. Consistent with this, aggressive mesenchymal GBM subtypes can acquire myeloid-associated transcriptional signatures associated with poor survival and resistance to radiotherapy and TMZ^42,45^. Thus, the interferon and MHC class II signatures induced by GSDME loss are compatible with an inflammatory mesenchymal state primed for antigen presentation and immune modulation^46^. We next classified the cells using the recently developed GBM framework of Nomura et al., which comprises 10 cellular states, including mesenchymal, hypoxia, and stress programs^3^. Across all four models, loss of GSDME consistently increased the components of these states **(Supplementary Figure 6G)**. Nevertheless, spatial profiling of the matched CME014 tumor revealed regional heterogeneity within the mesenchymal compartment, with podoplanin^high^ (PDPN^high^) and CD44^high^ tumor areas, corresponding to a GBM mesenchymal state as defined in ref^47^, showing lower GSDME expression than in adjacent PDPN^low^ CD44^low^-areas **(Figure 4I)**.

Unlike epithelial cancers, GBM does not follow a classical EMT framework but instead displays transitions between neural-like, progenitor-like, and mesenchymal-like states. In this context, mesenchymal programs are often viewed as adaptive differentiation or stress-associated states, whereas tumor-propagating capacity is linked to neurodevelopmental and progenitor-like programs^2,42,48^. The convergence of the transcriptional changes in GSDME-deficient cells resembled the injury response program, which is inversely related to a neurodevelopmental, progenitor-associated state in GBM stem cells^42^. Scoring these programs in our cells suggested that GSDME is involved in balancing these states **(Supplementary Figure. 6H)**. Consistent with this framework, GSDME deficiency suppressed glial progenitor-associated transcriptional programs **(Supplementary Figure 6A)**. Applying the Nomura et al. classification further showed that, specifically in CME037 and CME038, loss of GSDME was accompanied by a reduction in progenitor-like states **(Supplementary Figure. 7A)**. These findings prompt us to examine progenitor-associated tumor-propagating capacity.

GSDME expression positively correlated with CytoTRACE scores, suggestive of a less differentiated transcriptional state^49^ **(Supplementary Figure 7B)**, and was accompanied by increased surface abundance of CD49f, an integrin α-chain encoded by *ITGA6*, in CITE-seq data and previously linked to glioma stemness^50^ **(Figure 4J and Supplementary Figure 7C-E)**. In line with these observations, GSDME-deficient cells showed reduced CD133 levels **(Figure 4K)** and reduced neurosphere-forming frequency in limiting dilution assays **(Figure 4L and Supplementary Figure 7F)**. GSDME-deficient neurospheres also grew less than GSDME-proficient controls in 3D cultures **(Figure 4M)**. As in the GSDME KO models of CME037 and CME038, GSDME deficiency also reduced Developmental program scores in the two mesenchymal models **(Supplementary Figure 7G)**.

Notably, nuclear localization of GSDME was observed in GSDME^high^ tumor cells in human GBM tissue **(Figure 1C)** and overlap between GSDME expression and the Chromatin-Reg malignant state, a state enriched for chromatin and transcriptional regulators, was observed in two independent single cell transcriptomic cohorts **(Supplementary Figure 7H-K)**, raising the possibility that a nuclear pool of full-length GSDME might contribute to the transcriptional state dynamics identified here. To further investigate this, we classified GBM cells in the COMET cohort according to nuclear GSDME staining and found that the two classical tumors exhibited markedly increased nuclear GSDME localization than the mesenchymal tumor **(Figure 4N)**. Consistent with this pattern, nuclear GSDME^+^ GBM cells displayed significantly lower PDPN expression, suggesting that nuclear accumulation of full-length GSDME may be linked to attenuation of the mesenchymal transcriptional program **(Figure 4O)**.

To determine whether the functional effects of GSDME deficiency extend across species, we examined the murine glioma cell line GL261 and observed comparable phenotypic changes **(Supplementary Figure 7L-O)**, supporting a conserved role for GSDME in human and murine glioma models.

Together, these data support a cell autonomous role of full-length GSDME in actively reshaping transcriptional heterogeneity and stem-like features in GBM.

### GSDME tunes macrophage-induced phenotypic transitions

We next asked whether GSDME also shapes how GBM cells respond to macrophage-derived cues. TAMs have been shown to promote mesenchymal transitions in GBM through paracrine oncostatin M (OSM)-OSMR signaling^47^, and we detected OSM in supernatants from CME037 and CME038 co-cultured with THP-1-macrophages **(Supplementary Figure 8A)**. However, our data also showed that macrophage-derived S100A4 induces GSDME, which restrains MES-Hyp programs. We therefore investigated whether GSDME determines the type of mesenchymal state induced in the presence of macrophages.

We first revisited scRNA-seq co-culture datasets of CME037 and CME038 that included human monocyte-derived macrophages^30^. In both PDCL models, MES-Hyp abundance was reduced upon co-culture despite upregulation of other mesenchymal programs **(Supplementary Figure 8B)**. We then co-incubated GSDME-proficient and GSDME-deficient GBM cells with THP-1-macrophages and performed bulk RNA-seq on purified tumor cells from co-cultures and matched monocultures **(Supplementary Figure 8C)**. Principal component analysis (PCA) revealed four distinct clusters, indicating condition-specific transcriptional states **(Figure 5A)**. Gene set variation analysis (GSVA) further showed upregulation of EMT-related signatures upon macrophage co-culture **(Supplementary Figure 8D)**, confirming macrophage-induced mesenchymal remodeling.

**Figure 5.**
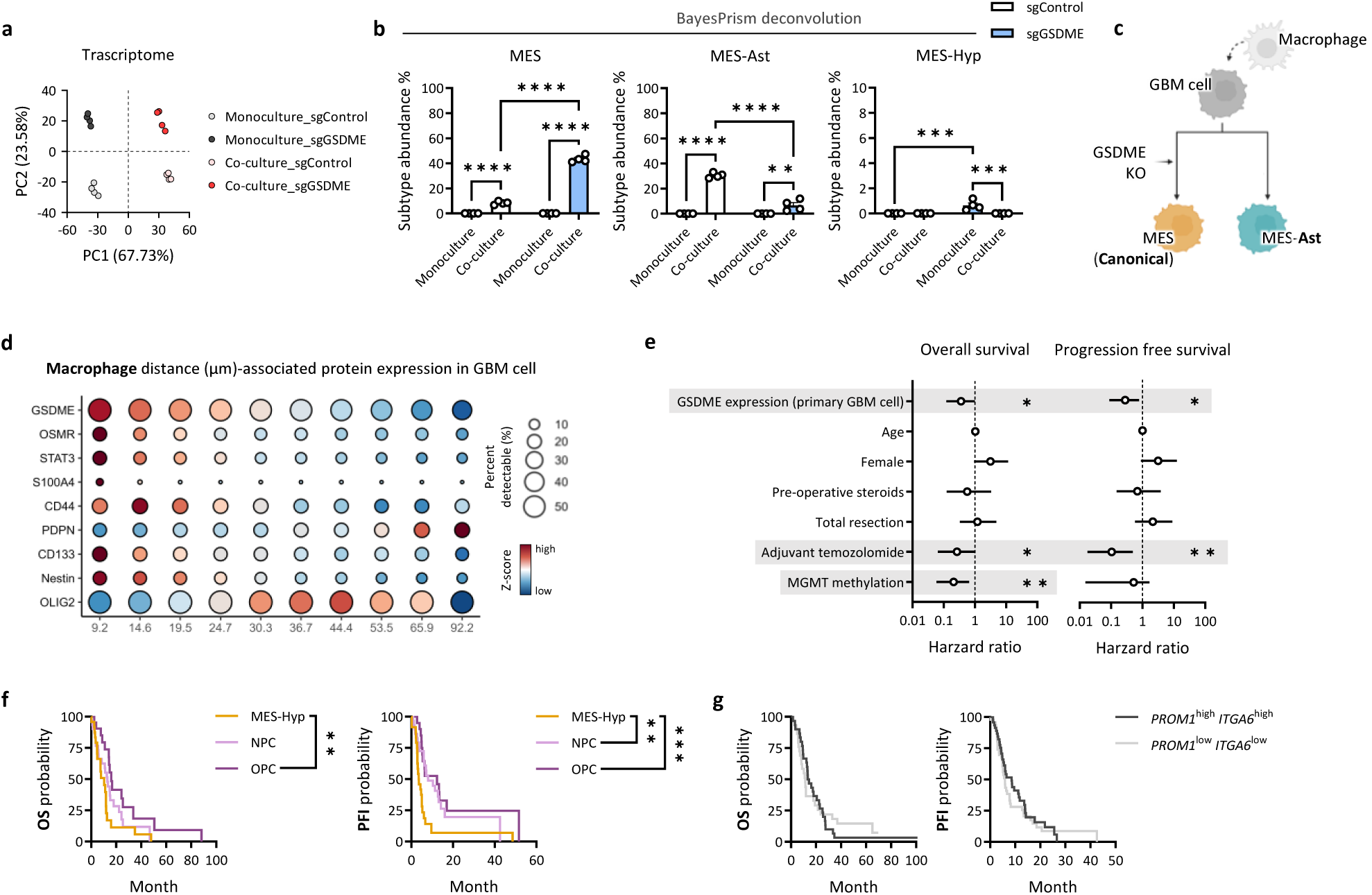
GSDME modulates macrophage-induced phenotypic transitions in GBM cells. **(A)** PCA of transcriptomic profiles from sgControl and sgGSDME PDCLs under monoculture and THP-1-macrophage co-culture conditions. **(B)** BayesPrism deconvolution of Greenwald et al. mesenchymal subtypes, showing the percentage of each subtype across the indicated culture and genotype conditions. **(C)** Schematic of GBM cell subtype transitions under macrophage interaction in GSDME-high or GSDME-low states. **(D)** Standardized protein expression per GBM cell across distance deciles using macrophages as index cells. **(E)** Multivariate Cox regression analysis of GSDME expression per GBM cell for OS and PFS, adjusted for clinical parameters. **(F)** OS and PFS analyses of TCGA GBM samples classified as MES-Hyp, NPC, or OPC. **(G)** OS and PFS analyses of TCGA GBM samples stratified as *PROM1*^high^ *ITGA6*^high^ or *PROM1*^low^ *ITGA6*^low^. Data are shown as mean ± SEM. Each dot in the bar plots represents an individual replicate. Statistical significance is indicated as *p < 0.05, **p < 0.01, ***p < 0.001, ****p < 0.0001.

Deconvolution of bulk RNA-seq data showed that macrophage exposure increased mesenchymal programs in both GSDME-proficient and GSDME-deficient cells, but with distinct subtype biases. In GSDME-proficient cells, macrophage-derived signals strongly induced MES-Ast programs, with more limited induction of canonical MES-like programs and no increase in MES-Hyp abundance **(Figure 5B)**. In line with this, differential expression analysis identified *SLC6A11*, an astrocyte-associated gene and component of the MES-Ast program, as upregulated under this condition **(Supplementary Figure 8E)**. By contrast, GSDME-deficient cells showed a stronger macrophage-induced increase in MES-like programs but a blunted MES-Ast response **(Figure 5B, left and middle)**. Although GSDME loss increased MES-Hyp under monoculture conditions, macrophage co-culture suppressed this MES-Hyp bias **(Figure 5B, right)**. Together, these findings suggest that GSDME does not simply determine whether macrophages induce mesenchymal remodeling but instead controls which mesenchymal state is adopted in response to macrophage-derived cues **(Figure 5C)**.

We next resolved these relationships spatially in tumor tissue with our COMET data. GBM cells proximal to macrophages showed higher GSDME and CD133 expression, together with increased expression of components of the mesenchymal-associated OSMR-STAT3 axis, whereas PDPN expression was lowest in these macrophage-proximal cells **(Figure 5D)**. Similar spatial gradients were observed around endothelial cells, although over a more restricted distance **(Supplementary Figure 8F)**, suggesting that the macrophage-associated pattern extends over a broader range and positioning GSDME within this spatially organized GBM cell state response.

Finally, we examined whether these GSDME-associated states were linked to clinical outcomes by integrating the protein staining with the available survival data from our MILAN cohort. High GSDME expression in GBM cells independently predicted improved overall survival (OS) and progression-free survival (PFS) in multivariable Cox regression models that included adjuvant TMZ treatment and MGMT promoter methylation as significant covariates **(Figure 5E)**. Consistent with these findings, analysis of the published CARE Consortium scRNA-seq cohort independently confirmed that higher *GSDME* expression in GBM cells was associated with delayed disease progression **(Supplementary Figure 8G)**.

Thus, although GSDME^high^ tumor cells are embedded within macrophage-rich regions, GSDME expression is associated with a state that restrains hypoxia-biased mesenchymal transition, providing a potential explanation for its favorable prognostic association. Consistently, TCGA GBM analyses using the Greenwald et al. framework^26^ showed shorter OS and PFS in patients with MES-Hyp-enriched tumors **(Figure 5F)**, in line with prior work linking MES-hypoxia abundance to worse prognosis in GBM^51^. Expression of the stemness-associated genes *PROM1* encoding CD133, and *ITGA6*, encoding CD49f, either alone or in combination, was not associated with OS or PFS **(Figure 5G, Supplementary Figure 8H and 8I)**.

### Coordination of GSDME-inflammasome-NINJ1 lytic program promotes tumor rejection

GSDME is a pore-forming executor of secondary necrosis and pyroptosis, suggesting that GSDME^high^ glioma cells are primed for inflammatory cell death under therapeutic stress. To test this, we compared TMZ, a weak inducer of immunogenicity, with canonical inducers of immunogenic cell death (ICD), including mitoxantrone (MTX)^52^, and hypericin-based photodynamic therapy (PDT), a ROS-inducing treatment previously shown to trigger robust tumor-rejecting responses in preclinical glioma models^53^. TMZ and MTX failed to induce a significant loss of viability, particularly in the recurrent PDCL **(Supplementary Figure 9A)**. In contrast, PDT curbed cell viability across all PDCLs in a dose-dependent manner, irrespective of the GBM stage or phenotypes **(Figure 6A and Supplementary Figure 9F)**. PDT-treated cells exhibited a faint but detectable caspase-dependent accumulation of the p30-NT pore-forming fragment of GSDME **(Figure 6B, 6C, Supplementary Figure 9B, 9G and 9H)**, along with pyroptotic ballooning morphology **(Figure 6D)**, consistent with a pre-lytic stage coupled to progressive Cytotox Green uptake via GSDME-mediated pores^19,54^. PDT also induced LDH release, indicating plasma membrane rupture (PMR) and cell lysis **(Figure 6E, Supplementary Figure 9D, 9I and 9J)**. In line with our previous studies^53,55,56^, PDT-induced cell death was accompanied by early ATP release followed by extracellular HMGB1 efflux, both established ICD-associated damage-associated molecular patterns (DAMPs) **(Figure 6E, Supplementary Figure 9E, 9I and 9J)**. GSDME deficiency did not affect the overall kinetics of cell death, as measured by proportion of Cytotox Green^+^ cells, nor did it alter LDH release or HMGB1 efflux **(Figure 6F, left, Supplementary Figure 9K, 9L, left and 9M)**. However, GSDME loss attenuated pyroptotic-like ballooning morphology **(Figure 6F, middle, right and Supplementary Figure 9L, right)** and early ATP release **(Figure 6G)**. This is congruent with the role of GSDMs in mediating membrane permeability changes, leading to swelling and osmotic imbalance uncoupled from full lysis^57,58^. Similar findings were obtained in murine GL261 cells, which constitutively express GSDME **(Supplementary Figure 9N)**. PDT induced robust cell death **(Supplementary Figure 9O)**, pyroptotic ballooning **(Supplementary Figure 9P)**, marked LDH release at 24 h **(Supplementary Figure 9Q)**, and DAMPs release **(Supplementary Figure 9R)**. GSDME KO likewise did not prevent PDT-induced cell death **(Supplementary Figure 9S)**, despite the loss of full-length GSDME and the consequent absence of PDT-induced GSDME-NT generation **(Supplementary Figure 9T)**.

**Figure 6.**
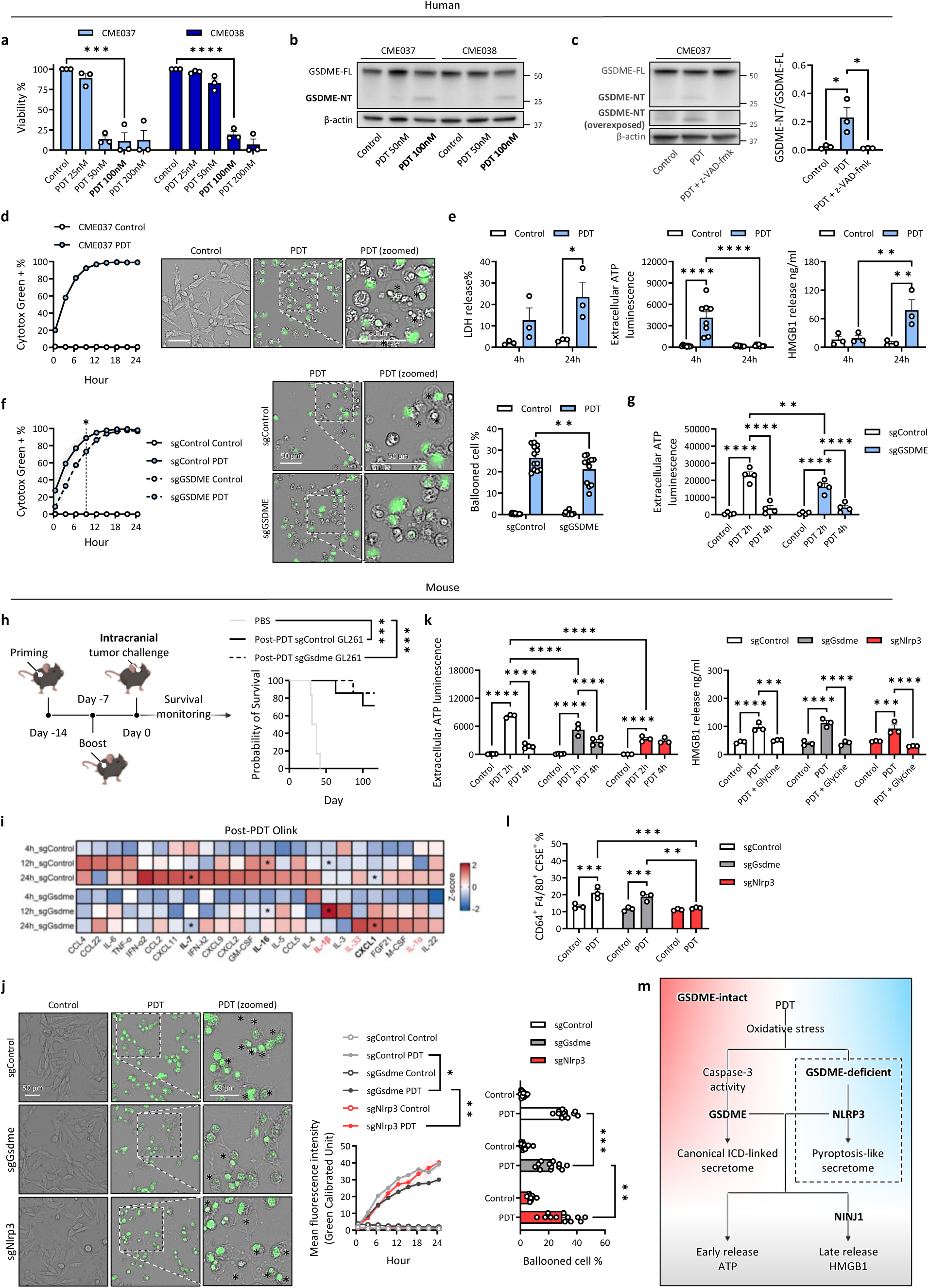
PDT activates a cooperative GSDME-inflammasome-NINJ1 lytic program that promotes tumor rejection. **(A)** Cell viability of CME037 and CME038 treated with increasing doses of Hyp-PDT, measured by CellTiter-Glo 2.0. **(B)** Immunoblot of full-length GSDME cleavage to the N-terminal fragment in CME037 and CME038 following Hyp-PDT. **(C)** Immunoblot and quantification of GSDME cleavage in CME037 following Hyp-PDT in the presence or absence of the pan-caspase inhibitor z-VAD-fmk. **(D)** Time-lapse quantification of Cytotox Green^+^ cells over 24 h (left) and representative morphology at 24 h (right) in CME037 following Hyp-PDT, acquired using the Incucyte live-cell imaging system. **(E)** LDH release, extracellular ATP levels, and HMGB1 release at 4 and 24 h in CME037 following Hyp-PDT. **(F)** Time-lapse analysis of Cytotox Green positivity in sgControl and sgGSDME CME037 after Hyp-PDT, including the percentage of Cytotox Green^+^ cells over time (left), representative late stage morphology (middle), and quantification of ballooned cell abundance (right). **(G)** Extracellular ATP release at early time points after Hyp-PDT in sgControl and sgGSDME CME037. **(H)** Schematic of the prophylactic tumor vaccination experiment using PDT-treated sgControl or sgGsdme GL261 (left) and Kaplan-Meier survival analysis (right). **(I)** Olink-based secretome profiling of sgControl and sgGsdme GL261 after Hyp-PDT at early, intermediate, and terminal time points (4, 12, and 24h, respectively). **(J)** Extracellular ATP release at early time points (left) and HMGB1 release at 24 h in the presence or absence of glycine (right) after Hyp-PDT in sgControl, sgGsdme and sgNlrp3 GL261 cells. **(K)** BMDM phagocytosis assay measuring engulfment of dying or dead sgControl, sgGsdme, and sgNlrp3 GL261 cells after Hyp-PDT. **(L)** Morphologic and kinetic analysis of lytic cell death in sgControl, sgGsdme and sgNlrp3 GL261 after Hyp-PDT, including representative 24-h morphology with or without Hyp-PDT (*, ballooned cells; left), time-lapse quantification of Cytotox Green mean fluorescence intensity (middle), and quantification of ballooned-cell abundance across genotypes (right). **(M)** Schematic of model in which Hyp-PDT activates a cooperative GSDME, inflammasome and NINJ1-dependent lytic death program in GBM cells. Data are shown as mean ± SEM. Each dot in the bar plots represents an individual replicate. Statistical significance is indicated as *p < 0.05, **p < 0.01, ***p < 0.001, ****p < 0.0001.

To validate whether these GSDME-mediated effects altered ICD efficacy in vivo, we performed prophylactic vaccination, the gold-standard criterion of ICD induction, as used in our previous study^56^. Mice were vaccinated, or not, subcutaneously with dying GSDME-proficient or GSDME-deficient GL261 cells, then rechallenged intracranially one week later with live GL261 cells **(Figure 6H, left)**. Remarkably, whereas all non-vaccinated mice developed brain tumors and showed rapid survival decline, both PDT-based vaccines elicited strong tumor rejection, with 70-80% of mice remaining tumor-free **(Figure. 6H, right)**. These findings suggest that the prominent tumor-rejecting ability of PDT-generated vaccines, likely relies on multiple signaling mechanisms, which may compensate for the lack of GSDME. To probe compensatory mechanisms supporting immunogenicity in the absence of GSDME, we profiled the GL261 cell secretome over time following PDT^59^, focusing on GSDM’s pore-mediated release of proinflammatory cytokines and chemokines as inducible DAMPs^18,24,60,61^. GSDME deficiency reduced the release of canonical ICD-associated cytokines^56,62^, including IL-6 and TNF, the caspase-3-processed cytokine IL-16^63^, and chemokines including CCL2, CCL5, CXCL9, and CXCL11 **(Figure 6I)**. By contrast, IL-1β and IL-33 secretion increased in GSDME-deficient cells **(Figure 6I)**. These findings suggest that GSDME loss skews the inflammatory secretome toward an inflammasome-dominant output, hinting to a contribution of NLRP3-mediated signaling^64^, which is likely caused by PDT induced ROS and ER stress^65–67^.

To test this directly, we generated NLRP3 KO cells in a GSDME-proficient background **(Supplementary Figure 9U)**. NLRP3 deficiency did not affect ballooning morphology or Cytotox Green uptake **(Figure 6J)**, but blunted the early ATP release **(Figure 6K, left)**, without altering HMGB1 and LDH efflux **(Figure 6K, right and Supplementary Figure 9V)**, and impaired macrophage engulfment of dying cells, consistent with a role of ATP as an early “find-me” signal in efferocytosis **(Figure 6L)**^68^. These data suggest that, following PDT, GSDME pore formation is the predominant driver of osmotic imbalance and the pre-lytic release of ATP, cytokines, and chemokines, while NLRP3 signaling facilitates early ATP release and phagocytic licensing.

Neither GSDME nor NLRP3 deficiency reduced HMGB1 release **(Figure 6K, right)**, indicating that the efflux of large molecular weight DAMPs is not primarily controlled by these pathways. Oligomerization of Ninjurin-1 (NINJ1), triggered by initial osmotic imbalance drives PMR, enabling the release of large proinflammatory molecules, such as LDH and HMGB1, the latter acting as a key danger signal driving glioma rejection in prophylactic DC-vaccination settings^58,69,70^. In line with this, glycine, which protects cells from the loss of osmolarity^71^, prevented both HMGB1 and LDH release in PDT-treated cells **(Figure 6K, right and Supplementary Figure 9V)**, without altering the development of ballooned morphology and Cytotoxic Green influx **(Supplementary Figure 9X and 9Y)** as reported^58^. Consistent with this, CME014 human GBM cells, which express low levels of NINJ1, showed attenuated HMGB1 and LDH release after PDT **(Supplementary Figure 9I and 9W)**.

Together these findings unravel that while GSDME governs the pre-lytic composition of the inflammatory secretome, in ROS-induced ICD, additional NLRP3-mediated signaling and NINJ1-dependent membrane rupture support the overall immunogenicity.

### GSDME-NINJ1 proinflammatory death module localizes to macrophage-rich perivascular niches in GBM

Given the complementary roles of caspase-dependent GSDME cleavage in pre-lytic pore formation and NINJ1-mediated PMR in terminal lysis, we next asked whether these components are co-expressed across cancer contexts and retained within the GBM architecture. Analysis of CCLE cancer cell line transcriptomes revealed a positive correlation between *GSDME* and *NINJ1* expression, with GBM cell lines among those exhibiting high expression of both genes **(Figure 7A)**. Consistently, GBM was the tumor lineage most enriched for *GSDME*/*NINJ1* double-high cell lines, showing the highest odds ratio across cancer types **(Figure 7B)**. In the MILAN cohort, GSDME protein expression was higher in primary than recurrent GBM tumor cells **(Figure 7C)**. To validate this pattern in an independent dataset, we revisited the publicly available scRNA-seq cohort comprising primary and recurrent GBM samples^72^ **(Supplementary Figure 7H)**. This confirmed reduced *GSDME* expression at recurrence, whereas *NINJ1* was increased in recurrent tumors **(Figure 7D)**. Thus, despite reduced *GSDME* expression, increased *NINJ1* expression suggests that the molecular machinery required for PDT-induced inflammatory lytic cell death is preserved after recurrence **(Figure 7E)**. We then asked whether these effectors were spatially organized within macrophage-rich, vascularized niches in GBM tissue. Spatial transcriptomic analysis revealed significant co-expression of GSDME, CASP3, and NINJ1 in macrophage-enriched perivascular regions **(Figure 7F and 7G)**. We explored whether this niche-associated lytic program could also be resolved across clinically defined tumor compartments. Analysis of a publicly available scRNA-seq dataset generated from MRI-guided regional sampling of GBM revealed enrichment of the lytic program consisting in *CASP3, GSDME, NINJ1* expression, in tumor core samples relative to peritumoral edema **(Supplementary Figure 10A and 10B)**. Together, these findings indicate that macrophage-rich GBM niches retain the molecular architecture required for ROS-induced GSDME-dependent pre-lytic permeabilization and NINJ1-dependent terminal DAMP release, with this lytic program preferentially enriched in radiographically defined tumor core regions relative to peritumoral edema.

**Figure 7.**
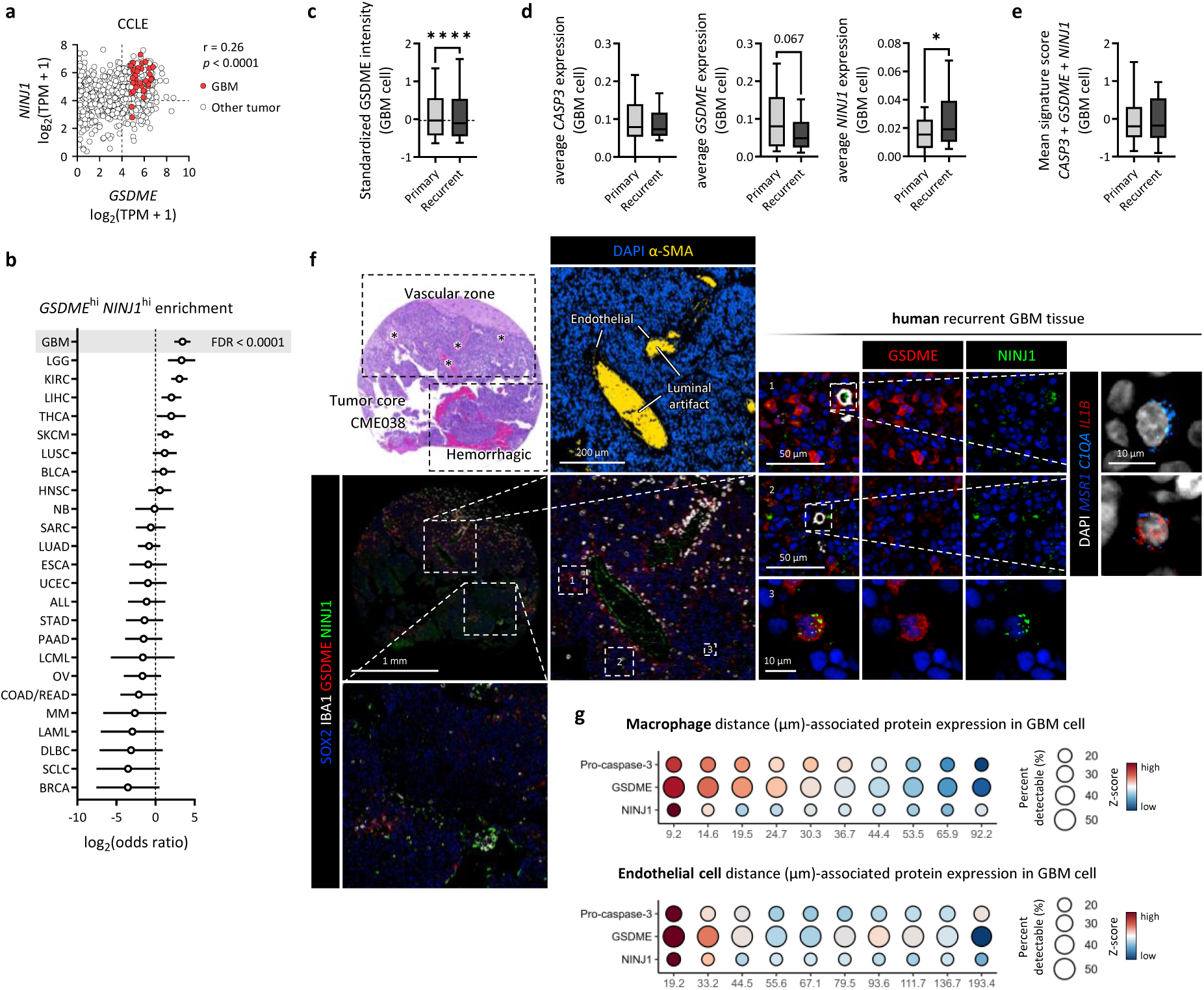
A proinflammatory death module localizes to macrophage-rich perivascular niches in human GBM. **(A)** Correlation between *GSDME* and *NINJ1* expression across pan-cancer cell lines from the CCLE dataset. **(B)** Odds ratio analysis of *GSDME*^hi^*NINJ1*^hi^ enrichment across pan-cancer cell lines. **(C)** Standardized GSDME expression per GBM cell in primary versus recurrent GBM samples from the MILAN cohort. **(D)** Box plots of *CASP3*, *GSDME* and *NINJ1* expression per GBM cell from primary versus recurrent GBM samples. **(E)** Hyp-PDT signature score in primary versus recurrent GBM. **(F)** Representative H&E image of the tumor-core region from the tumor corresponding to CME038, together with multiplexed COMET™ staining for SOX2, IBA1, GSDME, NINJ1 and α-SMA and RNAscope detection of *MSR1*, *C1QA* and *IL1B*. **(G)** Expression of caspase-3, GSDME and NINJ1 in GBM cells binned by distance deciles from macrophages or endothelial cells as index cells. Box plots show the median, interquartile range, and 10^th^-90^th^ percentile whiskers. Statistical significance is indicated as *p < 0.05, ****p < 0.0001.

## Discussion

GBM develops through reciprocal interactions with tumor-associated myeloid cells, whereby tumor-derived growth factors and metabolic and inflammatory cues recruit and reprogram myeloid cells, which in turn reinforce malignant adaptation and invasion^30,47,73^. However, how this crosstalk is spatially organized, how it regulates protein expression in cancer cells and how these niche-dependent programs influence cancer cell states and therapeutic vulnerabilities is incompletely defined. Here, by integrating spatial and single-cell multi-omics across primary and recurrent GBM with functional experiments, we identify GSDME as a niche-instructed determinant of GBM-macrophage crosstalk that links macrophage-associated malignant cell states to ICD potential.

Spatial analysis across primary and recurrent GBM identified macrophage-associated regions in which malignant cells exhibited elevated GSDME expression. This relationship was most prominent around infiltrating macrophages and was less evident around resident microglia, indicating that tumor cell GSDME is associated with a specific myeloid context. The proximal macrophages displayed scavenger-like and immunosuppressive features, and GSDME^high^ malignant cells were frequently organized within macrophage-rich perivascular regions. Notably, GSDME remained predominantly full length in these areas, with little evidence of caspase-3 activation, indicating that GSDME expression marked a viable malignant cell state rather than ongoing lytic cell death. S100A4 neutralization and inhibition of EGFR-Sp1 signaling further supported a pathway through which suppressive macrophages regulate GSDME expression in adjacent tumor cells.

Under non-lytic conditions, loss of GSDME shifted GBM cells toward canonical and hypoxia-associated mesenchymal programs. GSDME also influenced the state adopted in response to macrophage-derived cues. Whereas macrophage exposure favored a MES-Ast program in GSDME-proficient cells, GSDME-deficient cells preferentially acquired a canonical MES-like program. These findings extend previous work on macrophage-derived OSM-OSMR signaling^47^ by showing that macrophage-induced mesenchymal remodeling is not a uniform response but is shaped by the molecular state of the receiving tumor cell. GSDME therefore biases mesenchymal state selection rather than simply preventing mesenchymal transition.

High tumor-cell GSDME was associated with longer survival and delayed progression in the examined cohorts. This association is consistent with the restriction of MES-Hyp programs, which have been linked to invasion, treatment resistance, recurrence, and T cell dysfunction^26,29,72,74^. However, because GSDME also supported progenitor- and stem-associated features, it should not be interpreted as a simple tumor suppressor. Rather, it influences the balance of malignant cells between progenitor-like and distinct mesenchymal states. This context-dependent redistribution may help explain why GSDME expression within an immunosuppressive macrophage niche is nevertheless associated with more favorable clinical outcomes.

Our findings suggests GSDME-FL engages nuclear regulatory mechanisms in GBM. Its nuclear localization raises the possibility of interactions with transcriptional or chromatin-associated regulators. During preparation of this manuscript, Du et al. reported that nuclear GSDME associates with SMARCA5 and NCL to increase chromatin accessibility at PLK4 promoter, thereby promoting melanoma progression^75^. This independently supports a non-pyroptotic nuclear function for GSDME, but the phenotypic outputs vary between tumor types. Whereas GSDME promotes epithelial-to-mesenchymal transition in melanoma, GSDME in GBM restricts specific mesenchymal programs. How GSDME accumulates in the nucleus of tumor cells, whether GSDME engages similar chromatin-regulatory machinery in GBM and which molecular partners mediate this function, remain to be determined.

Importantly, this GSDME^high^ state in viable cells creates a latent inflammatory cell death potential that an appropriate therapeutic stress can unleash. We found that in contrast to standard-of-care chemotherapy (i.e. TMZ) or other immunogenic chemotherapy (i.e. MTX), assessed in this study, or the caspase activator raptinal^25^, hypericin-based PDT engages the full potential of proinflammatory cell death in both primary and recurrent GBM. PDT, a potent ROS-inducing treatment, elicited robust tumor rejection in prophylactic vaccination assays, consistent with previous evidence that PDT can generate effective DC-based anti-glioma vaccines^53^.

Mechanistically, we uncovered a multilayered death program driven by PDT where GSDME, inflammasome signaling and NINJ1 cooperate complementarily, with each module contributing to distinct signaling functions that collectively maintain high tumor rejecting ability in vivo. GSDME promoted rapid osmotic swelling and the pre-lytic release of ICD-associated cytokines and chemokines, whereas NLRP3 signaling contributed to early ATP efflux and enhanced myeloid phagocytic engagement. Terminal release of lysis-associated DAMPs was instead mediated by a glycine-sensitive, likely NINJ1-dependent PMR step. These DAMPs included HMGB1, which is required for robust PDT-induced DC-based anti-glioma immunity and may promote DC immunogenicity by facilitating the sensing of extracellular DNA^53,76^.

The spatial co-expression of GSDME and NINJ1 within macrophage-rich and vascularized niches indicates that molecular components capable of supporting inflammatory cell death are retained within the human GBM architecture. This may be particularly relevant at recurrence, when therapeutic options are limited and immune checkpoint blockade has largely failed^77,78^. Whether this pattern can serve as a predictive spatial biomarker for responses to PDT^79^ or clinically relevant ICD inducers, including radiotherapy, remains to be established. Prospective studies should also determine whether spatially targeted sampling can preserve or reconstitute this macrophage-associated state ex vivo for personalized vaccine development^80–83^.

Together, our findings show how cellular context governs GSDME output. Reciprocal GBM-macrophage crosstalk sustains a predominantly full-length GSDME pool that biases tumor cell plasticity, whereas proteolytic activation enables GSDME to participate in a coordinated inflammatory cell death program. This framework expands GSDME biology beyond pore formation and places it at the intersection of spatial niche signaling, tumor cell identity and inflammatory cell death.

## Methods

### Human GBM specimens

#### COMET and RNAscope cohort

Human GBM formalin-fixed paraffin-embedded (FFPE) specimens were retrieved under protocol S62248, approved by the Ethics Committee of UZ Leuven. Written informed consent was obtained where required, and specimens were pseudonymized before transfer and analysis. All tumors were obtained during routine surgical resection. Recurrent GBM specimens were obtained at reoperation for tumor recurrence. The COMET cohort consisted of FFPE specimens from two patients with IDH^wt^ GBM. Matched primary and recurrent GBM specimens from one patient are referred to throughout the study as CME037 and CME038, respectively, and were represented by 10 and 5 tissue cores. A primary GBM specimen from a second patient is referred to as CME014 and was represented by 1 tissue core. These specimen names correspond to the matched patient-derived GBM cell lines (PDCLs) generated from the same tumors.

#### MILAN spatial proteomics cohort

Primary and recurrent human GBM FFPE samples were retrieved from the previously described retrospective GBM cohort under protocols S59804, S61081, S62248, 18/0021R, 19/0021R and METC 16-4-022, approved by the medical ethics review boards of the participating hospitals^84^. Written informed consent was obtained where required, and specimens were pseudonymized before transfer and analysis. All specimens were obtained during routine surgical resection. Recurrent GBM specimens were obtained at reoperation for tumor recurrence. Available treatment annotations are summarized with the clinical characteristics in Supplementary Figure. 1j. The MILAN dataset comprised 202 arrayed FFPE tissue cores. Of these, 195 cores were linked to study-level identifiers corresponding to 33 patients. The remaining 7 cores were not linked to a study-level patient identifier and were included in tissue-level analysis only.

### Experimental models

#### Patient-derived GBM cell lines

PDCLs were obtained from the laboratory of Frederik De Smet under protocols S61081 and S59804, approved by the Ethics Committee of UZ Leuven. Written informed consent was obtained where required, and all patient-derived material was pseudonymized before transfer and analysis. Cells were maintained as serum-free adherent cultures on laminin-coated flasks in DMEM/F12 (ThermoFisher, 11320074) with an in-house neural stem cell supplement, antibiotic-antimycotic (Sigma-Aldrich, A5955), heparin (STEMCELL, 7980), EGF (STEMCELL, 78006) and FGF2 (STEMCELL, 78003). The in-house supplement, adapted from a previously published formulation for serum-free neural stem cell culture^85^, contained glucose, HEPES (Gibco, 12509079), L-glutamine (Thermo Scientific, 25030024), apo-transferrin (Gibco, 11108016), insulin (Sigma-Aldrich, I2643), putrescine, sodium selenite and progesterone. EGF and FGF2 were each used at a final concentration of 20 ng/mL. Cultures were maintained at 37°C and 5% CO₂. For passaging and experimental plating, cells were detached with Accutase (Gibco, 11599686). Cultures were routinely tested for mycoplasma contamination.

#### THP-1 cells and THP-1-derived macrophages

THP-1 monocytic cells were maintained from laboratory stock in RPMI 1640 medium (Gibco, 61870036) supplemented with 10% FBS and 1% antibiotic-antimycotic at 37°C and 5% CO₂. Cells were cultured in suspension and routinely tested for mycoplasma contamination. For macrophage differentiation, THP-1 cells were seeded at 1.5 × 10⁵ cells per well in 6-well plates and treated with 100 nM phorbol 12-myristate 13-acetate (PMA; MedChemExpress, HY-18739) for 48 h. Differentiated cells were washed and rested in PMA-free complete medium for 24 h before downstream experiments. Where indicated, THP-1-derived macrophages (THP-1-macrophages) were polarized before co-culture, conditioned-medium collection, or molecular analysis. Polarization was induced by treating cells for 24 h with either IFN-γ (20 ng/mL; STEMCELL, 78020.1) and LPS (100 ng/mL; Sigma-Aldrich, L2630) or IL-4 (20 ng/mL; STEMCELL, 78045.1).

#### HMC3 microglial cells

HMC3 cells were provided by the laboratory of Gabriele Bergers (VIB-KU Leuven Center for Cancer Biology) and maintained in DMEM/F12 medium supplemented with 10% FBS, 1% antibiotic-antimycotic, and 1% L-glutamine at 37°C and 5% CO₂. Cells were passaged using TrypLE and routinely tested for mycoplasma contamination.

#### GL261 murine glioma cells

GL261 cells were maintained from laboratory stock in DMEM supplemented with 10% FBS and 1% antibiotic-antimycotic at 37°C and 5% CO₂. Cells were passaged using 0.25% Trypsin-EDTA and routinely tested for mycoplasma contamination. For in vivo tumor rechallenge experiments, GL261 cells were harvested on the day of intracranial inoculation and prepared as single-cell suspensions before injection.

#### Murine bone marrow-derived macrophages

Murine bone marrow-derived macrophages (BMDMs) were generated from bone marrow cells isolated from 6-8 weeks old female C57BL/6J mice. Bone marrow cells were cultured in non-tissue-culture-treated 10 cm Petri dishes in DMEM supplemented with 20% FBS and 30% L929-conditioned medium. Cells were initially cultured in 6 mL medium, and an additional 3 mL was added on day 3. On day 7, adherent macrophages were detached with ice-cold PBS and used for downstream experiments.

#### Mouse models

All animal experiments were approved by the KU Leuven Animal Ethics Committee under P19/2020 and performed in accordance with the Guide for the Care and Use of Laboratory Animals: Eighth Edition2, the Oncology Best practices: Signs, Endpoints and Refinements for in Vivo Experiments (OBSERVE) guidelines3, EU Directive 2010/63/EU, Regulation (EU) 2019/1010 and Belgian national legislation. Female C57BL/6JOlaHsd mice (Inotiv) were housed under standard conditions at the KU Leuven Imaging and Pathology: MicroImaging (IPMI) facility. Mice were 10 weeks old at the start of the experimental procedures.

### Integrated COMET spatial proteomics and RNAscope in situ hybridization

#### COMET and RNAscope staining and image acquisition

FFPE GBM tissue microarrays were profiled on the COMET platform (Lunaphore Technologies) by combining RNAscope HiPlex Pro in situ hybridization (Advanced Cell Diagnostics, Inc.) with sequential immunofluorescence (seqIF) on the same tissue sections^86^. Tissue microarrays were generated from 2 mm punches taken from FFPE GBM tissue blocks using the TMA Grand Master system (3DHistech). Sections were cut at 3 µm, mounted on supercharged Flex IHC microscope slides (K8020, Dako/Agilent), and baked for 1 h at 52°C. Slides were deparaffinized on an Autostainer XL system (Leica) using three xylene baths followed by 100% ethanol and 70% ethanol incubations. Heat-induced epitope retrieval was performed in a PT Module instrument (Epredia) at 99°C for 1 h using Dewax and HIER Buffer H, pH 9 (AR03, Lunaphore Technologies). Slides were briefly washed in Multistaining Buffer (BU06, Lunaphore Technologies) before loading onto the COMET instrument.

RNA detection was performed using the RNAscope HiPlex Pro for COMET Reagent Kit (322075, Advanced Cell Diagnostics, Inc.). Sections were incubated with RNAscope HiPlex Pretreat Pro for 30 min at 40°C and hybridized with the RNAscope HiPlex Pro probe pool for 2 h at 40°C. Signal amplification was performed with RNAscope HiPlex Pro AMP 1, AMP 2, and AMP 3 reagents for 30 min each at 37°C. Cyclic signal detection and cleavage were performed using RNAscope HiPlex Pro Cleaving Reagent diluted in 4× saline-sodium citrate buffer for 15 min at 27°C. Three detection-cleavage cycles were performed to resolve the eight-plex RNA panel. The same sections were then processed by sequential immunofluorescence on the COMET platform to detect a 32-plex protein panel. For each cycle, Alexa Fluor Plus secondary antibodies (Thermo Fisher Scientific) were applied as a mixture of two species-complementary antibodies together with DAPI. Images were acquired using fixed exposure times: DAPI, 25 ms; TRITC, 250 ms; and Cy5, 400 ms. The assay generated multilayer OME-TIFF files containing DAPI, autofluorescence reference channels, and target-specific RNA and protein fluorescence channels.

Representative COMET and RNAscope images shown in the Figure.s were generated in HORIZON Viewer (Lunaphore Technologies) after channel-specific background subtraction, with display settings adjusted uniformly across compared images.

#### Image preprocessing, segmentation and single cell feature extraction

Stained images were processed before single cell analysis by subtracting the autofluorescence signal measured in the corresponding unstained baseline scan acquired prior to marker staining. Individual cellular objects were segmented using StarDist^87^, and morphological and marker intensity features were extracted for each segmented cell. Marker intensities were normalized by z-score transformation, with z-scores trimmed to −5 to 5 to reduce the influence of outliers^88^. Cell identities were assigned using CONCLAVE for lineage and structural markers^89^, followed by threshold-based annotation of selected functional marker states. Annotation quality was assessed by the marker expression dot plot across cell types. Single cell-resolved marker intensities were used to compare the expression of selected markers between cell types and to perform downstream spatial analysis.

#### Nearest neighbor analysis and community analysis

To quantify the relationship between GBM cell protein expression and spatial context, Euclidean distances were calculated separately for each tissue sample and annotated region from each GBM cell to the nearest cell of each specified neighboring population. The analysis were limited to the tumor core and infiltrative margin, and the peripheral region was excluded. For each region and neighboring population, samples were included if they contained at least 30 GBM cells and 30 cells of the neighboring population. Models were fitted when at least 50 GBM cell observations were available across a minimum of five samples. Distances were log-transformed and standardized before modeling. For each protein, GBM cell marker intensity was modeled as a function of distance to the specified neighboring population, with sample included as a random effect to account for within-sample variability. *P* values were adjusted across all tests using the Benjamini-Hochberg method. For volcano plots, adjusted *P* values were displayed as -log_10_(FDR). Negative coefficients indicate higher protein expression in GBM cells closer to the specified neighboring population, whereas positive coefficients indicate higher expression at greater distances.

To visualize distance-associated expression patterns, GBM cells were ranked by their distance to the nearest cell of the indicated neighboring population within each sample and region and divided into proximity deciles, with each decile containing approximately equal numbers of GBM cells. For each proximity decile, mean marker intensity and the percentage of marker-positive GBM cells were calculated for each sample and then averaged across samples for visualization. Mean expression values were z-score normalized across proximity deciles and displayed as dot plots.

Community analysis was performed using SCIMAP^90^ with a neighborhood radius of 200 pixels, corresponding to approximately 56 μm.

### MILAN spatial proteomics

#### MILAN staining and image acquisition

Multiplex spatial proteomic analysis was performed using the Multiple Iterative Labeling by Antibody Neodeposition (MILAN) platform on FFPE tissue sections. Following deparaffinization and antigen retrieval, tissue sections underwent iterative rounds of immunofluorescence staining. During each cycle, a subset of antibodies together with DAPI nuclear counterstain was applied, followed by fluorescence image acquisition. After imaging, antibody complexes were chemically dissociated from the tissue while preserving antigen integrity, enabling subsequent staining cycles on the same section. This process was repeated until the complete marker panel had been acquired. Whole-slide fluorescence images were generated after each staining round using a high-resolution fluorescence imaging platform.

#### MILAN image preprocessing, segmentation and single cell feature extraction

Raw MILAN scans were converted into grayscale 16-bit TIFF images using ZEN software. Images from sequential staining cycles were registered using the Python package imreg. Stained images were processed before single-cell analysis by subtracting the autofluorescence image acquired before staining from the measured signal of the corresponding tissue section. Individual cellular objects were segmented using StarDist^87^, and morphological and marker-intensity features were extracted for each segmented cell. Marker intensities were normalized by z-score transformation, with z-scores trimmed to a range of −5 to 5 to reduce the influence of outliers^88^. Cell identities were assigned using CONCLAVE^89^ and threshold-based annotation for SOX2, TMEM119, CD163, and CD68. Annotation quality was assessed by principal component analysis (PCA) of normalized marker intensities and by the marker expression dot plot across annotated cell types.

#### Quantitative analysis of MILAN data

For MILAN image level analysis, GSDME was quantified at single cell resolution within annotated GBM cells. For each tissue core, normalized GSDME intensities from individual GBM cells were summarized as mean GBM cell GSDME intensity and paired with the relative abundance of annotated macrophages or microglia in the same core. Cores were included in the correlation analysis only if GBM cells represented at least 25% of all annotated cells, to restrict the analysis to tumor-containing cores with sufficient malignant cells. Associations between GBM cell GSDME intensity and myeloid abundance were assessed using Pearson correlation at the tissue core level.

To assess spatial proximity, Euclidean distances were calculated within each tissue core from every GBM cell to the nearest cell of each indicated myeloid population. GBM cells were ranked by distance to the indicated myeloid population and divided into proximity deciles, with each decile containing approximately equal numbers of GBM cells (Figure. 1f). For each proximity decile, GBM cell GSDME intensity was summarized and visualized as a distance-associated trend plot. Differences of GSDME expression across proximity deciles were assessed by one-way ANOVA. For clinical association analysis, GSDME intensity was quantified from single cell-resolved measurements within annotated GBM cells. Tissue cores containing fewer than 10 GBM cells were excluded. For each remaining tissue core, mean normalized GBM cell GSDME intensity was calculated and aggregated at the patient level using the number of GBM cells per core as weights. Patients with fewer than 30 total GBM cells after filtering were excluded from survival analysis. Associations between patient-level GBM-cell GSDME intensity and overall survival (OS) or progression-free survival (PFS) were assessed using Cox proportional hazards regression.

### Public single cell and bulk RNA-seq analysis

#### GBM PDCL scRNA-seq analysis

Public GBM patient-derived cell lines (PDCLs) scRNA-seq data was analyzed from downloaded count matrix and metadata. Analysis was performed in R using a Seurat-based workflow^91^. Count matrix and metadata were used to generate Seurat objects, followed by ambient RNA correction with decontX^92^ and doublets removal with scDblFinder^93^. Cells were normalized, scaled, and embedded by PCA and uniform manifold approximation and projection (UMAP). Subtype signature scores were calculated for each cell and used to assign transcriptional states, enabling reconstruction of the subtype composition of selected PDCLs.

#### GBM-macrophage co-culture scRNA-seq analysis

Public GBM-macrophage co-culture scRNA-seq data (EGAS00001007482) was analyzed together with publicly available monoculture scRNA-seq data from PDCLs. Analysis was performed in R using a Seurat-based workflow^91^. Count matrix and metadata were used to generate Seurat objects, followed by ambient RNA correction with decontX^92^ and doublets removal with scDblFinder^93^. Cells were normalized, scaled, and embedded by PCA and uniform manifold approximation and projection (UMAP).

Published cell type and sample annotations were retained. For comparison of monoculture and macrophage co-culture conditions, the analysis was restricted to PDCLs represented in both datasets: CME037, CME038, and LBT003. *GSDME* expression was evaluated within the GBM cell population, visualized using UMAP-based Nebulosa density plots and box plots, and compared between culture conditions using the Wilcoxon rank-sum test.

For cell-cell communication analysis, macrophages and GBM cells from the corresponding co-cultures were analyzed. GBM cells were stratified by *GSDME* detectability, with detectable and undetectable *GSDME* expression defining *GSDME*^high^ and *GSDME*^low^ receiver populations, respectively. Macrophage-to-GBM cell ligand-receptor interactions were inferred using LIANA for each receiver context^33^. Ligand-receptor pairs identified in the macrophage-to-*GSDME*^high^ context were compared with those identified in the macrophage-to-*GSDME*^low^ context, and pairs unique to the *GSDME*^high^ context were prioritized using LIANA aggregate ranking.

For CME037 and CME038, GBM transcriptional states of PDCLs in each culture condition were assigned using the Greenwald et al. classification framework^26^. Cell state proportions were calculated within each PDCL model and culture condition for visualization.

#### GBM tissue scRNA-seq analysis

Public GBM tissue scRNA-seq datasets (GSE84465, GSE117891 and GSE174554) were analyzed from downloaded count matrix and metadata. Datasets were processed in R using the Seurat-based workflow described above, including ambient RNA correction, doublet removal, normalization, scaling, PCA, and UMAP visualization. Where required, batch effects were corrected using Harmony^94^, and Harmony-corrected dimensions were used for nearest-neighbor graph construction, clustering, and visualization.

Cell clusters were identified using CHOIR^95^, and broad cell type annotation was performed using SingleR^96^. Malignant cells were inferred using SCEVAN^97^. For each dataset, malignant GBM cells were extracted to compare *GSDME* expression across the indicated conditions. *GSDME* expression was visualized using UMAP-based Nebulosa density plots and compared across groups using the Wilcoxon rank-sum test.

Myeloid cells were extracted separately to assess macrophage and microglial state composition. Myeloid subtypes were assigned using the Miller et al. myeloid framework^32^, and subtype proportions were calculated within each sample before comparison across conditions. Changes in myeloid subtype composition were visualized as indicated in the corresponding Figure.s.

#### Bulk RNA-seq analysis

Public GBM bulk RNA-seq data (GSE136751) was analyzed from raw count matrix. Counts were normalized as counts per million (CPM) and log-transformed. *GSDME* and *GSDMD* expression were compared across the indicated conditions using an unpaired t test.

#### TCGA data analysis

TCGA GBM RNA-seq expression data and curated survival annotations were obtained from UCSC Xena. Expression values were downloaded as log_2_(TPM + 1)-transformed values, and survival annotations included overall survival (OS) and progression free interval (PFI). Primary GBM samples with matched expression and survival data were retained for analysis. Gene Set Variation Analysis (GSVA) scores were calculated using signatures from the Greenwald et al. GBM classification framework^26^. Samples with dominant MES-Hyp, NPC, or OPC states were used for state level survival analysis. For gene level survival analysis, expression values for *PROM1* and *ITGA6*, encoding CD133 and CD49f respectively, were extracted. Upper- and lower-quartile groups were defined for each gene. For combined marker analysis, samples classified as upper quartile for both *PROM1* and *ITGA6* were defined as dual high, whereas samples classified as lower quartile for both genes were defined as dual low. OS and PFI were analyzed using Kaplan-Meier curves, and survival differences were assessed by log-rank test.

#### CCLE data analysis

Cancer Cell Line Encyclopedia (CCLE) pan-cancer RNA-seq expression data and cell line metadata were obtained from the DepMap portal. Expression values were downloaded as transcripts per million (TPM) and transformed as log_2_(TPM + 1). The correlation between GSDME and NINJ1 expression across cell lines was assessed using Pearson’s correlation coefficient. For each tumor type, enrichment of cell lines with concurrent high GSDME and NINJ1 expression was quantified by calculating odds ratios. Confidence intervals were estimated for each odds ratio, and *P* values were adjusted for multiple testing using the false discovery rate method.

#### GL261 scRNA-seq analysis

Public GL261 cells scRNA-seq dataset (GSE244301) was analyzed from downloaded count matrix and metadata. Datasets were processed in R using the Seurat-based workflow described above, including ambient RNA correction, doublet removal, normalization, scaling, PCA, and UMAP visualization. Where required, batch effects were corrected, and the corrected dimensions were used for nearest-neighbor graph construction, clustering, and visualization. *Gsdme* expression was visualized using UMAP-based Nebulosa density plots. MES-Hyp, MES, and MES-Ast signatures from the Greenwald et al. human GBM classification framework^26^ were converted to murine gene symbols before signature scoring. Signature scores were visualized using Nebulosa density plots. CytoTRACE2 was used to estimate relative differentiation state^49^, and CytoTRACE2 scores were visualized on the UMAP embedding.

### Co-culture of GBM PDCLs and myeloid lineage cells

PDCLs were seeded on laminin-coated 6-well plates in serum-free GBM culture medium before co-culture. CME037 and CME038 cells were seeded at 6 × 10⁵ cells per well, whereas CME014 cells were seeded at 1.5 × 10⁵ cells per well, reflecting differences in cell size and growth morphology. At the start of co-culture, PDCL cultures were approximately 70% - 80% confluent.

Co-cultures were performed using 0.4 μm pore-size cell culture inserts to maintain indirect contact between PDCLs and myeloid cells. PDCLs were cultured in the lower chamber, and THP-1 cells, THP-1-macrophages, polarized THP-1-macrophages or HMC3 cells were seeded in the upper insert chamber (Figure. 2a,h and Supplementary Figure. 2e,f). This configuration allowed exchange of soluble factors while preventing migration of co-cultured cells between chambers. Co-cultures were maintained for 4 days in 2 mL medium per well. PDCL monocultures were processed in parallel as controls.

After 4 days of co-culture, inserts containing myeloid cells were removed, and PDCLs in the lower chamber were washed and harvested for downstream assays. Where required, co-culture supernatants were collected before cell harvest, cleared by centrifugation to remove residual cells and debris and used for soluble-factor analysis.

### S100A4 neutralization and pathway inhibition in GBM-macrophage co-cultures

For S100A4 neutralization, PDCLs were co-cultured with THP-1-macrophages using the indirect co-culture system described above. S100A4 neutralizing antibody clone 6B12 (V3S-0622-YC2540, Creative Biolabs) was added to the co-culture medium at a final concentration of 6 μg/mL. To maintain antibody exposure at the intended final concentration during the co-culture period of 4 days without substantially changing the culture volume, 6B12 was replenished daily from day 2 to day 4 by adding 50 μL antibody-containing medium to each well. For pathway inhibition, PDCLs were co-cultured with THP-1-macrophages and treated with inhibitors during the co-culture period. EGFR signaling was inhibited with erlotinib at 0.5, 1 or 2 μM. JAK-STAT signaling was inhibited with ruxolitinib at 100, 150 or 200 nM. Sp1-dependent transcriptional activity was inhibited with plicamycin at the indicated concentration. Untreated THP-1-macrophage co-cultures and matched PDCL monocultures were processed in parallel as controls. At endpoint, inserts containing THP-1-macrophages were removed, and PDCLs were harvested for downstream assays.

### Reverse transcription quantitative PCR

Total RNA was extracted from PDCL monocultures and PDCLs harvested after indirect co-culture using TRIzol™ reagent according to the manufacturer’s protocol. RNA concentration and purity were assessed by NanoDrop. cDNA was synthesized using the QuantiTect Reverse Transcription Kit. Quantitative PCR was performed using SYBR Green chemistry in technical triplicate on a 7500 Fast Real-Time PCR System. Relative transcript abundance was calculated using the ΔΔCt method and normalized to 18S.

### Immunoblotting

Cells were lysed in RIPA buffer supplemented with protease and phosphatase inhibitors. Lysates were cleared by centrifugation, and protein concentration was determined using a BCA assay. Protein lysates, 50 μg per sample, were resolved by SDS-PAGE and transferred to membranes. Membranes were processed using either conventional blocking in 5% skim milk in TBS-T or Blotting Accelerator (Proteintech), according to the manufacturer’s instructions. Primary antibodies were incubated overnight at 4°C unless processed with Blotting Accelerator, followed by HRP-conjugated secondary antibodies. Primary antibodies used for immunoblotting targeted GSDME, GSDME N-terminal fragment (GSDME-NT), GSDMD, EGFR, phosphorylated EGFR, STAT3, phosphorylated STAT3, EGFRvIII, NINJ1, NLRP3, CD133, FRA1 and β-actin as indicated in the corresponding figures. Chemiluminescent signals were detected using ECL, and band intensities were quantified using Image Studio Lite.

### ELISA

Co-culture supernatants were collected before cell harvest and cleared by centrifugation to remove residual cells and debris. S100A4 concentrations were measured using the Human S100A4 ELISA Kit (Abcam) according to the manufacturer’s instructions. Absorbance values were converted to S100A4 concentrations using assay-specific standard curves. Supernatants from THP-1-macrophage monocultures and matched PDCL monocultures were processed in parallel as controls.

### nELISA secretome profiling

Co-culture supernatants were collected before cell harvest and cleared by centrifugation to remove residual cells and debris. Clarified supernatants were submitted to Nomic Bio for multiplex soluble-protein profiling by nELISA, a bead-based multiplex immunoassay platform described previously^98^. Amphiregulin and oncostatin M measurements were extracted from the normalized nELISA matrix and compared across culture conditions.

### CRISPR-Cas9-mediated gene knockout

GSDME knockout (KO) PDCLs and GSDME or NLRP3 knockout GL261 cells were generated by CRISPR-Cas9 ribonucleoprotein nucleofection using the 4D-Nucleofector system (Lonza). Synthetic sgRNAs targeting human *GSDME* or murine *Gsdme* or *Nlrp3* were complexed with recombinant Cas9 protein and delivered using the corresponding Lonza nucleofection solution and program. After nucleofection, cells were recovered in pre-warmed culture medium and expanded under the corresponding culture conditions for each model. Knockout efficiency was assessed by immunoblotting for the corresponding target protein. Control cells were processed in parallel using the corresponding nucleofection control condition.

### In-house GSDME KO PDCL scRNA-seq

#### GSDME KO scRNA-seq library generation and preprocessing

scRNA-seq libraries were generated from sgControl and sgGSDME PDCLs using the Chromium Next GEM Single Cell 3′ v3 workflow. CME037, CME038, CME014 and CME016 were profiled under both conditions. Single cell suspensions were counted and assessed for viability before labeling with sample-specific hashtag antibodies. Labeled cells were pooled by condition into sgControl and sgGSDME libraries and loaded for single cell capture. Libraries were sequenced on an Illumina sequencing platform. Raw sequencing data were processed with the 10x Genomics Cell Ranger pipeline to generate filtered gene expression matrix and hashtag count matrix. Hashtag counts were used to assign cells to PDCL identities.

#### GSDME KO scRNA-seq data analysis

Gene expression matrix, hashtag matrix, and sample annotations were analyzed in R using the Seurat-based workflow described above. Seurat objects were generated by retaining genes detected in at least three cells and cells with at least 200 detected features. Cells with more than 20% mitochondrial transcripts were excluded, followed by doublets removal using scDblFinder. Cells were normalized, scaled, and embedded by PCA and UMAP. GBM transcriptional states were assigned using the Neftel et al.^2^ and Greenwald et al.^26^ classification frameworks. Cell state proportions were calculated within each PDCL model and genotype for visualization. Gene set enrichment analysis (GSEA) was performed using malignant cell metaprograms (MPs) to identify transcriptional programs associated with GSDME loss^39^. Expression of selected genes was visualized using dot plots. Differential expression between sgGSDME and sgControl cells was performed separately within each PDCL model using MAST^99^. PDCL level differential expression results were combined across models using Fisher’s method, followed by Benjamini-Hochberg correction. Consensus genes were defined based on concordant directionality across PDCL models. Developmental and injury response signatures from Richards et al. were scored using UCell^42,100^. Transcription factor regulon activity was inferred using pySCENIC^101^. *FOSL1* regulon activity was extracted from the regulon activity matrix and compared between sgControl and sgGSDME cells within CME037 and CME038 separately.

### In-house GSDME KO GBM-macrophage co-culture bulk RNA-seq

#### Co-culture bulk RNA-seq library generation and preprocessing

CME037 sgControl and sgGSDME PDCLs were cultured as monocultures or in indirect co-culture with THP-1-macrophages as described above. Four biological replicates were prepared for each condition. After co-culture, inserts containing THP-1-macrophages were removed, and PDCLs in the lower chamber were washed before RNA extraction. Total RNA was extracted using a Qiagen RNA extraction kit according to the manufacturer’s instructions. Strand-specific, poly(A)-enriched mRNA-seq libraries were generated from quality-controlled RNA and sequenced on an Illumina NovaSeq X Plus platform using paired-end 150-bp reads. Sequencing was performed to a target depth of approximately 30 million read pairs per sample. Reads were aligned to GRCh38/hg38, and gene level count matrix were generated for downstream analysis.

#### Co-culture bulk RNA-seq data analysis

Count matrixes were analyzed in R. Raw counts were used for DESeq2 modeling, and variance stabilized expression values were used for PCA. TPM and log_2_(TPM + 1)-transformed expression values were generated. Epithelial-mesenchymal transition (EMT) activity was quantified using GSVA with the Hallmark EMT signature. GBM state deconvolution was performed using BayesPrism^102^. Monoculture CME037 cells from the public scRNA-seq analysis described above were used as single cell reference, with Greenwald et al. GBM classification framework annotations as state label^26^. Inferred state compositions were compared across monoculture and THP-1-macrophage co-culture conditions. To identify genotype-dependent transcriptional responses to macrophage co-culture, raw counts were modeled in DESeq2 using biological replicate, genotype, culture condition, and the genotype-by-culture interaction term. The interaction coefficient was used to identify genes whose co-culture response differed between sgControl and sgGSDME PDCLs. Interaction log_2_FC were moderated using apeglm to stabilize effect size estimates, and significant interaction genes were visualized by the volcano plot.

### GBM PDCL CITE-seq

Gene expression, antibody-derived tag (ADT) and sample annotation matrices from six PDCL models were analyzed in R using Seurat. Seurat objects were generated by retaining genes detected in at least three cells and cells with at least 200 detected features, followed by doublet removal using scDblFinder. RNA and ADT data were processed separately and integrated using Seurat’s weighted-nearest-neighbor framework, followed by PCA and multimodal UMAP visualization. Associations between GSDME RNA expression and ADT abundance were assessed across all cells and within each PDCL model using Spearman correlation. Linear mixed-effects models incorporating PDCL identity as a random intercept were used to identify ADTs positively associated with GSDME expression, followed by Benjamini-Hochberg correction.

### Seahorse experiments

Mitochondrial respiration and glycolytic activity of sgControl and sgGSDME PDCLs were measured using a Seahorse XFe24 extracellular flux analyzer. PDCLs were seeded on laminin-coated Seahorse assay plates and maintained under standard PDCL culture conditions before analysis. On the day of measurement, culture medium was replaced with Seahorse XF assay medium supplemented with metabolic substrates, and cells were equilibrated in a non-CO_2_ incubator before analysis. Oxygen consumption rate (OCR) and extracellular acidification rate (ECAR) were measured at baseline and after sequential injection of the indicated metabolic modulators. OCR and ECAR values were normalized to total protein content assessed by BCA assay.

### Limiting dilution assay

sgControl and sgGSDME PDCLs were assessed by in vitro limiting dilution assay. Single cell suspensions from four paired PDCL models were seeded into flat bottom ultra-low attachment 96-well plates in PDCL culture medium described above. Cells were plated by row using a two-fold dilution series from 2000 cells per well to approximately one cell per well. Cultures were maintained for 14 days, after which wells were scored as positive or negative based on the presence of tumor-sphere-like cell clusters. Sphere-initiating cell frequency and statistical significance between sgControl and sgGSDME conditions were determined by extreme limiting dilution analysis using the ELDA implementation in the R package statmod.

### Tumor sphere growth assay

sgControl and sgGSDME PDCLs were assessed in the tumor sphere growth assay in 3D. Cells were seeded at 2000 cells per well in U-bottom ultra-low attachment 96-well plates in the PDCL culture medium, allowing formation of a single aggregate per well. Sphere growth was monitored by bright-field microscopy every 2-3 days, as indicated in the corresponding Figure.s, for a total culture period of 10 days. Sphere size was quantified from microscopy images using ImageJ, and equivalent diameter was calculated for each sphere. Growth dynamics between sgControl and sgGSDME conditions were compared by two-way ANOVA. sgControl and sgGsdme GL261 cells were assessed with the same protocol in the FBS-free culture medium.

### Photodynamic therapy treatment

Photodynamic therapy (PDT) was performed in PDCLs and GL261 cells using hypericin as the photosensitizer. Cells were seeded to approximately 85% confluency before treatment, with seeding density adjusted for each cell model. PDCLs were treated in phenol red-free PDCL medium, whereas GL261 cells were treated in serum-free medium. After 2 h of hypericin incubation, PDCLs were changed to fresh phenol red-free PDCL medium without hypericin before irradiation. For GL261 cells, hypericin-containing medium was removed and replaced with PBS. Cells were irradiated at a fluence of 3.24 J/cm². After irradiation, PDCLs were maintained in the same fresh hypericin-free phenol red-free PDCL medium, whereas GL261 cells were cultured in serum-containing medium. Where indicated, z-VAD-fmk was added at 50 μM to inhibit caspase activity. Cells or supernatants were collected at the indicated time points for downstream assays.

### Mitoxantrone and temozolomide treatment

PDCLs were treated with mitoxantrone (MTX) at 4 μM for 24 h or temozolomide (TMZ) at 250 μM for 7 days. Untreated cells were processed as controls. Cell viability was assessed using CellTiter-Glo 2.0.

### CellTiter-Glo 2.0 viability assay

Cell viability after TMZ, MTX, or PDT treatment was measured using the CellTiter-Glo 2.0 viability assay (Promega) according to the manufacturer’s instructions. At the indicated time points, CellTiter-Glo 2.0 reagent was added directly to each well, and luminescence was measured using a plate reader. Viability was normalized to the corresponding untreated control condition, as indicated in the corresponding Figure.s.

### Incucyte live-cell imaging

PDT-induced cell death was monitored using Cytotox Green staining and the Incucyte S3 live-cell imaging system (Sartorius). After irradiation, Cytotox Green was added at 250 nM according to the manufacturer’s instructions, and plates were transferred to the Incucyte S3. Bright-field and green fluorescence images were acquired every 3 h using a 20× objective. Cytotox Green signal was quantified using Incucyte S3 software. Analysis definitions were optimized for each cell model using the first biological replicate and then applied consistently across replicates from the same model. Mean green fluorescence intensity and the percentage of Cytotox Green-positive cells were extracted for quantification. Representative bright-field and fluorescence images were exported and processed in ImageJ for display. Late time-point bright-field and fluorescence images were exported for morphology-associated quantification as described below.

### Ballooned cell quantification

Bright-field and fluorescence images from Incucyte live-cell imaging were used for morphology-associated quantification. Segmentation masks generated by the Incucyte Cell-by-Cell module were exported with the corresponding calibrated images and analyzed in CellProfiler^103^. Masks were imported as primary objects, and per-cell morphological and texture features were extracted from bright field images. Fluorescence images were retained as matched channels. Morphological phenotypes were classified using CellProfiler Analyst with supervised machine learning^104^. Representative cells were manually annotated to train a random forest classifier, with 100-150 cells annotated per phenotype. Training images were anonymized for treatment condition during annotation. The trained classifier was applied to all segmented cells to assign phenotype labels, and the proportion of ballooned cells was calculated for each image.

### Extracellular ATP assay

Extracellular ATP release after PDT was measured using Adenosine 5′-triphosphate (ATP) assay mix (Sigma-Aldrich). Culture supernatants were collected at the indicated time points after treatment, centrifuged to remove residual cells and debris, and transferred to white 96-well plates. Samples were diluted in HEPES buffer and mixed with ATP assay reagent immediately before measurement. Luminescence was measured using a plate reader and quantified.

### HMGB1 release assay

HMGB1 release after PDT was measured using the Lumit HMGB1 Human/Mouse Immunoassay (Promega) according to the manufacturer’s instructions. Culture supernatants were collected at the indicated time points after treatment, centrifuged to remove residual cells and debris. Samples were incubated with the HMGB1 antibody mixture, followed by addition of Lumit detection reagent. Luminescence was measured using a plate reader, and HMGB1 release was quantified using a positive-control dilution series.

### LDH release assay

LDH release after PDT was measured using the CytoTox 96 Non-Radioactive Cytotoxicity Assay (Promega) according to the manufacturer’s instructions. Culture supernatants were collected at the indicated time points after treatment, centrifuged to remove residual cells and debris. Samples were incubated with CytoTox 96 reagent. Reactions were stopped with Stop Solution, and absorbance was measured at 490 nm. LDH release was calculated after background correction and normalized to the corresponding control conditions.

### Prophylactic vaccination and orthotopic GL261 challenge

For prophylactic vaccination, GL261 sgControl or sgGsdme cells were treated with PDT as described above to generate dying/dead tumor cells, with approximately 80% cell death before vaccination. PDT-treated cells were collected, washed once with PBS, and resuspended at 5 × 10^5^ cells in 200 μL PBS per vaccination dose. Mice received subcutaneous vaccination with PDT-treated GL261 cells, or PBS as a negative control, on day -14, followed by a booster injection on day -7 in the contralateral flank.

On day 0, mice were orthotopically challenged by intracranial injection of live GL261 cells. Mice were anesthetized with ketamine/xylazine, placed in a stereotactic frame, and injected in the right striatum using a Hamilton microsyringe. A total of 5 × 10^4^ GL261 cells in 2 μL culture medium was injected at 2.5 mm lateral from the midline, 0.5 mm anterior to bregma, and 2.5 mm depth. The incision was closed with sutures after injection.

Mice were monitored for body weight and neurological symptoms at least three times per week. Humane endpoints were defined by loss of 20% initial body weight or grade 3-4 neurological symptoms using an adapted clinical scoring system: grade 0, no symptoms; grade 1, mild hemiparesis; grade 2, moderate hemiparesis with unstable gait; grade 3, severe hemiparesis, circling behavior, frequent falls, inability to move, or hunched posture; and grade 4, moribund state. Survival was analyzed by Kaplan-Meier analysis.

### Olink secretome profiling

Cytokines in supernatants from PDT-treated GL261 cells of the indicated genotypes were quantified using the Olink Mouse Cytokine Target 48 panel according to the manufacturer’s instructions. Undiluted samples were incubated with cytokine-specific antibody probes, followed by probe amplification using a T100 Thermal Cycler (Bio-Rad) and quantitative PCR-based readout on the Olink Signature Q100 system. Data quality control and normalization were performed using Olink NPX Signature software (2.0.2.0).

### Murine BMDM phagocytosis assay

BMDMs were prepared as described above. GL261 sgControl or sgGsdme cells were treated with PDT or maintained as untreated controls. Two hours after light irradiation, GL261 cells were collected and labeled with CellTrace CFSE. CFSE-labeled GL261 cells were co-incubated with BMDMs at a 1:1 ratio for 2 h at 37°C and 5% CO_2_. After co-incubation, cells were washed with PBS, counted and incubated with Fc receptor blocking antibody before surface staining. Cells were stained with fixable viability dye and fluorophore-conjugated antibodies against CD64 and F4/80. Phagocytosis was quantified by flow cytometry as the fraction of live CD64^+^F4/80^+^ BMDMs positive for CFSE signal, indicating uptake of labeled glioma cells.

### Statistical analysis

Statistical analyses were performed using GraphPad Prism. Comparisons between two groups were performed using two-tailed unpaired Student’s t tests. Comparisons involving more than two groups were performed using one-way or two-way ANOVA. Comparisons of gene expression between groups in scRNA-seq datasets were performed using two-sided Wilcoxon rank-sum tests. Survival analysis were performed using the Kaplan-Meier method and compared using log-rank tests. Associations between GSDME expression, clinical parameters, and survival outcomes were evaluated using Cox proportional-hazards regression. Statistical significance thresholds, statistical tests, sample sizes, and definitions of error bars are provided in the corresponding figure legends.

## Acknowledgements

The authors thank Niels Vandamme and Janick Mathys from the VIB Single Cell Core and Prof. Stephanie Humblet-Baron from the Immunobiology Unit at the Rega Institute for technical support and access to infrastructure. We thank the VIB Tech Watch Core for providing a Technology Adoption Grant to support the nELISA experiments. We also thank Natalia Lysak, Marleen Derweduwe, Prof. Uwe Himmelreich, Shannon Helsper, Chiara Caprioli, Lien Solie, and Kristine Rillaerts for their expert technical support. P.A. is supported by the KU Leuven C1 Consortium InterAction (C14/21/095), grants from the Research Foundation–Flanders (FWO; grant nos. G0A3320N and G094922N), the EOS MetaNiche consortium (no. 40007532), the iBOF ATLANTIS consortium (iBOF/21/053), and Research project funded by Kom op tegen Kanker (Stand up to Cancer). Y.Y. is supported by the scholarship of KU Leuven. C.B.G.-B receives postdoctoral research fellowship from Stichting tegen Kanker (Foundation against Cancer).

## Author Contributions

Y.Y., S.D.V. and P.A. designed and interpreted the experiments. Y.Y performed and analyzed the majority of the experiments with the technical help of E.V. In vitro experiments were performed by Y.Y. and E.V. In vivo experiments were performed by J.V.N. and A.Z.Z. In silico analyses were performed by Y.Y. with the help of K.A.N. F.D.S. provided the patient-derived cell lines. Sample’s pathology was performed by R.S. Microscopy images were acquired and analyzed by Y.Y. Genetically modified cell lines were generated by Y.Y. and E.V. MILAN image acquisition was performed by N.D. and analysis was performed by Y.Y with the help of K.A.N. COMET image acquisition was performed by J.S.C. and N.D. and analysis was performed by Y.Y. with the help of K.A.N. Phagocytosis experiments were performed by Y.Y. and S.Z. M.M., K.J. and S.M. provided expertise and discussed results. Y.Y. used BioRender to create the schematics. Y.Y. and P.A. wrote and edited the manuscript. P.A. and S.D. conceptualized and P.A. directed the study. All authors discussed the results and commented on the manuscript.

---

**Supplementary Figure 1.**
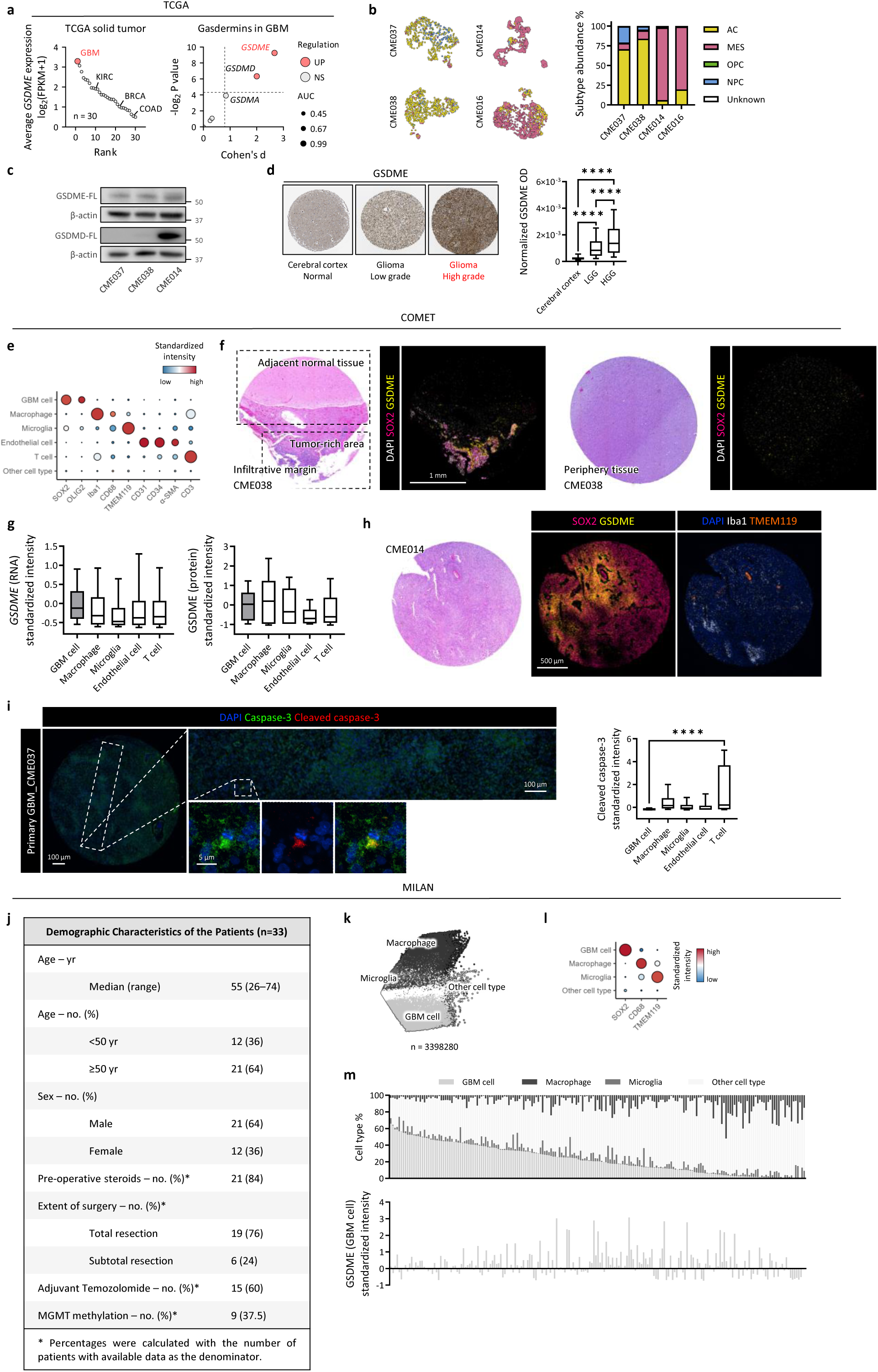
Full-length GSDME is elevated in GBM and enriched in tumor cells. **(A)** Pan-cancer ranking of mean GSDME expression across 30 solid tumor types from TCGA, ordered by decreasing GSDME expression (left). Effect sizes and statistical significance for tissue-level comparisons of gasdermin family member expression between GBM tumor tissue and normal brain tissue from TCGA are shown (right). **(B)** UMAP visualization of scRNA-seq profiles from 12 GBM PDCLs, including six matched primary-recurrent pairs. CME037/CME038 and CME014/CME016 represent two matched primary-recurrent pairs. Bar plots show the composition of Neftel et al. GBM cellular states in these four PDCLs. **(C)** Immunoblot analysis of GSDME and GSDMD expression in CME037, CME038, and CME014. **(D)** Representative immunohistochemistry images of GSDME in normal cortex, low-grade glioma, and high-grade glioma from HPA (left), with quantification of GSDME optical density (OD) across the full cohort (right). **(E)** Dot plot showing expression of markers used for cell type annotation in the COMET study. **(F)** COMET images of tumoral (left) and periphery (right) tissue corresponding to the PDCL CME038, showing SOX2, GSDME, and DAPI. **(G)** Single cell comparison of *GSDME* RNA expression measured by RNAscope (left) and GSDME protein intensity measured by COMET staining (right) across annotated cell types in the in-house COMET cohort. **(H)** COMET images of tumor tissue corresponding to the mesenchymal-dominant PDCL CME014, showing SOX2, GSDME, Iba1, TMEM119, and DAPI. **(I)** Representative COMET images of tumor tissue corresponding to PDCL CME037, showing total caspase-3, cleaved caspase-3, and DAPI (left), with single cell quantification of cleaved caspase-3 abundance across cell types in the full cohort (right). **(J)** Patient demographic and clinical characteristics for the MILAN study cohort. **(K)** PCA visualization of all segmented cells from the MILAN study, colored by annotated cell type. **(L)** Dot plot showing expression of markers used for cell type annotation in the MILAN study. **(M)** Bar plots showing cell type composition for each image (top) and GSDME intensity per GBM cell in the corresponding image (bottom) in the MILAN study. Box plots show the median, interquartile range, and 10^th^-90^th^ percentile whiskers. Statistical significance is indicated as ****p < 0.0001.

**Supplementary Figure 2.**
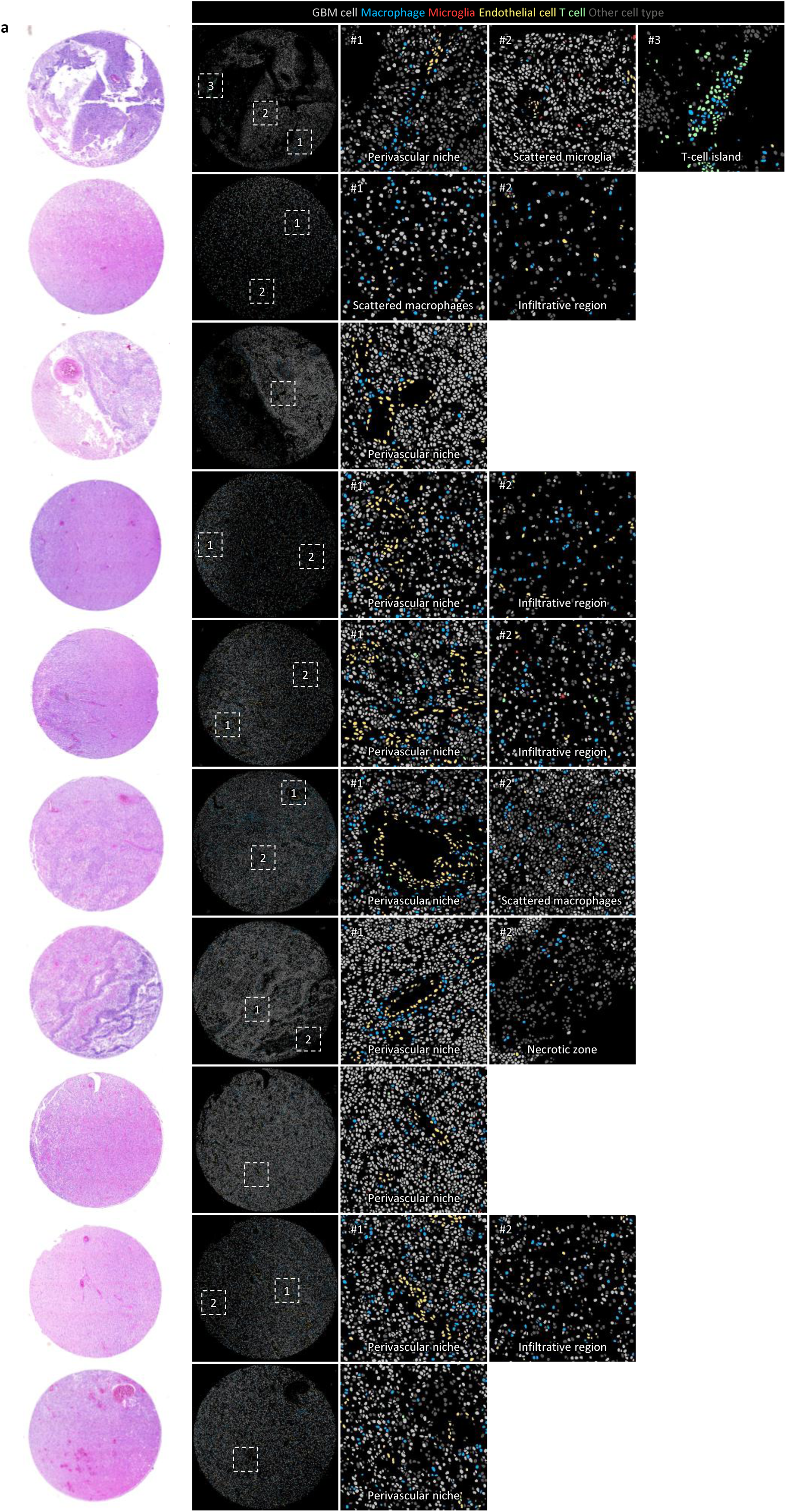
Histology and digital reconstruction of COMET samples corresponding to CME037. **(A)** H&E-stained tissue section, digital reconstruction of cell type annotations and magnified views of selected heterogeneous niches.

**Supplementary Figure 3.**
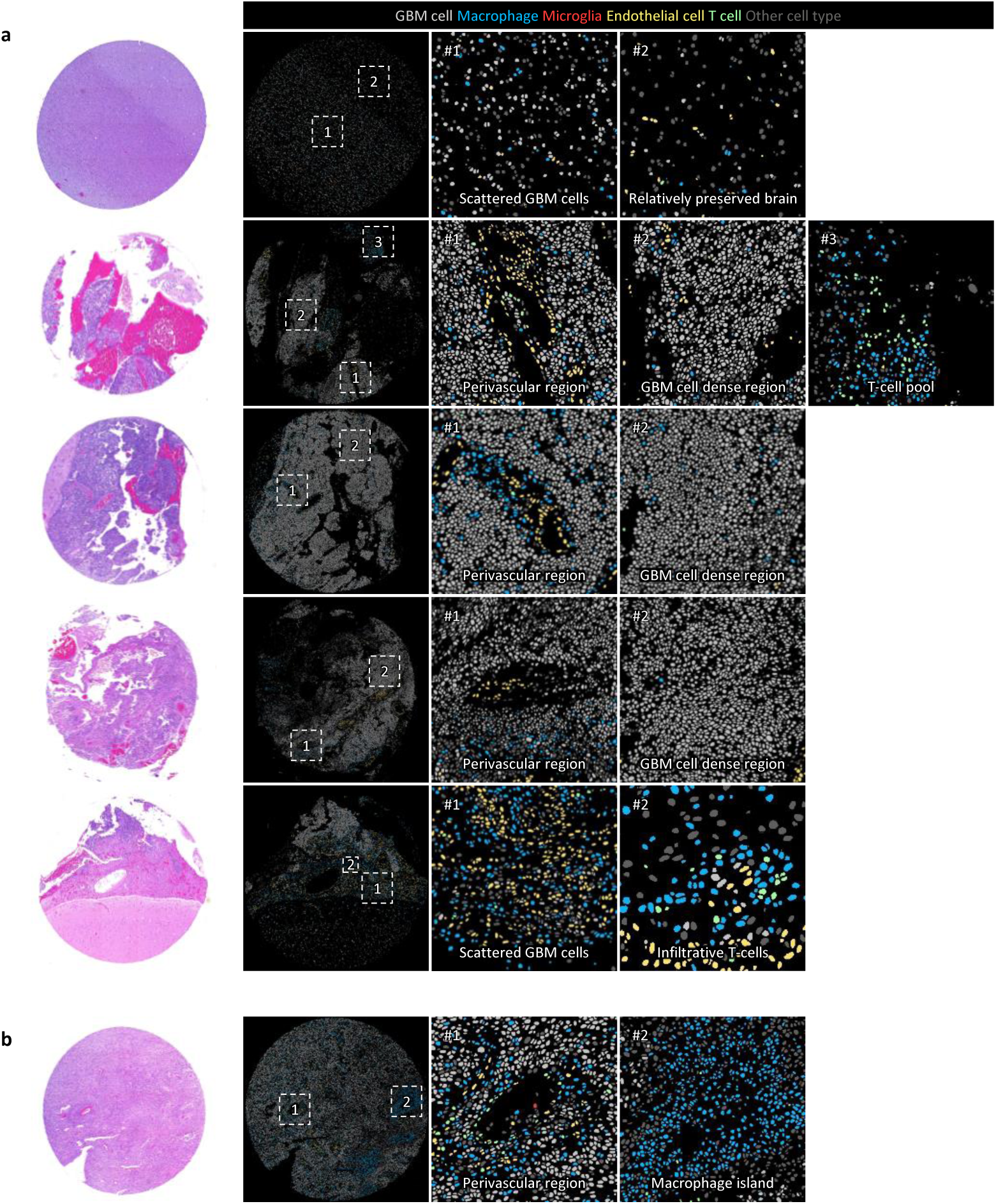
Histology and digital reconstruction of COMET samples corresponding to CME038 and CME014. **(A)** H&E-stained tissue section corresponding to CME038, digital reconstruction of cell type annotations and magnified views of selected heterogeneous niches. **(B)** H&E-stained tissue section corresponding to CME014, digital reconstruction of cell type annotations and magnified views of selected heterogeneous niches.

**Supplementary Figure 4.**
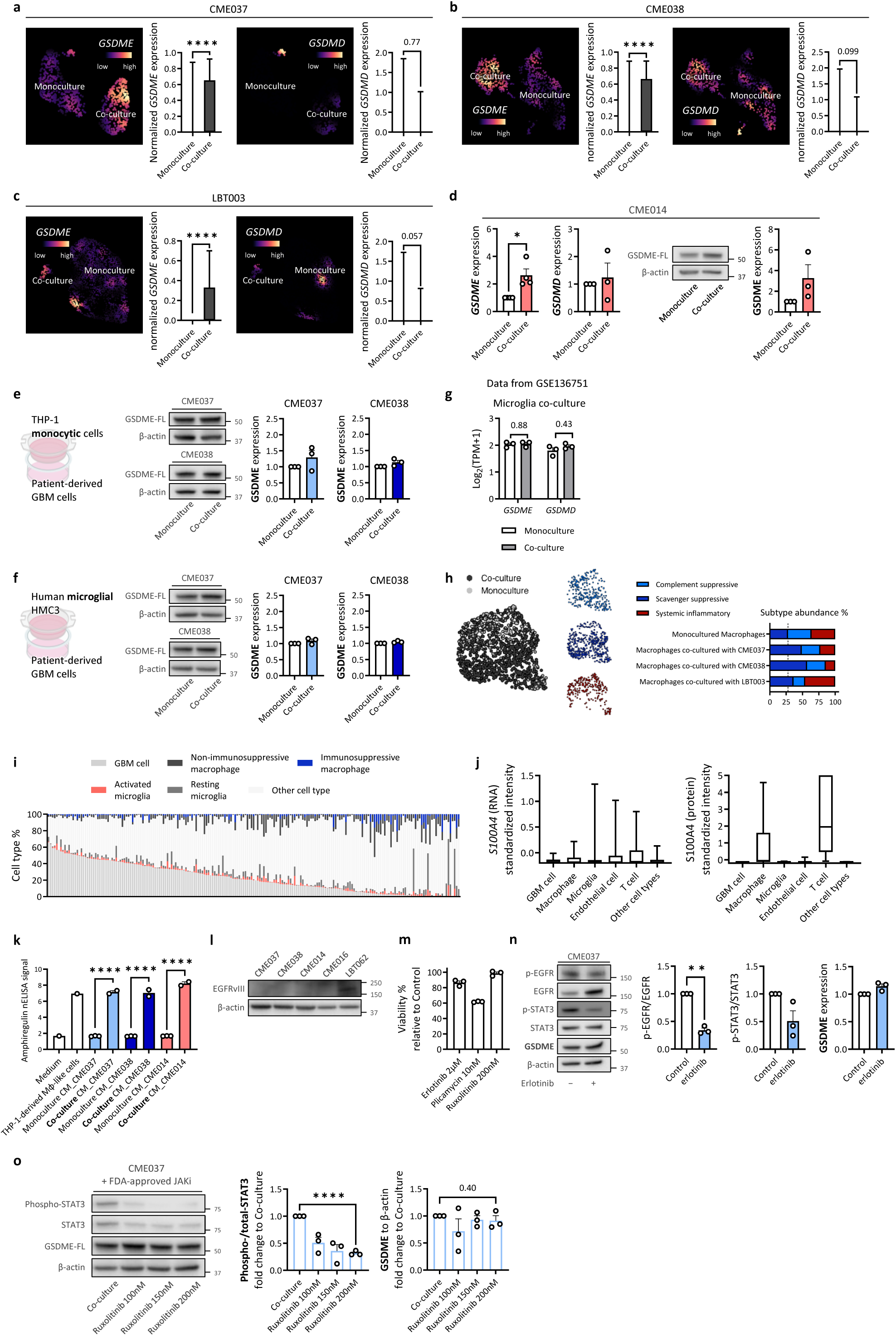
Macrophage co-culture selectively enhances GSDME expression in GBM cells. **(A)** Nebulosa density plots and box plots show *GSDME* and *GSDMD* expression of CME037 maintained in monoculture or co-cultured with human macrophages. **(B)** Nebulosa density plots and box plots show *GSDME* and *GSDMD* expression of CME038 maintained in monoculture or co-cultured with human macrophages. **(C)** Nebulosa density plots and box plots show *GSDME* and *GSDMD* expression of LBT003 maintained in monoculture or co-cultured with human macrophages. **(D)** qPCR analysis of *GSDME* and *GSDMD* expression and immunoblots of GSDME expression in CME014 maintained in monoculture or indirectly co-cultured with THP-1-derived macrophages. **(E)** Immunoblot analysis of GSDME expression in CME037 and CME038 maintained in monoculture or indirectly co-cultured with THP-1 monocytic cells. **(F)** Bulk RNA-seq analysis of GSE136751 comparing GSDME and GSDMD expression in GBM cells maintained in monoculture or co-cultured with microglia. **(G)** Immunoblot analysis of GSDME expression in CME037, CME038, and CME014 maintained in monoculture or indirectly co-cultured with MHC3 microglia. **(H)** UMAP visualization of scRNA-seq profiles from macrophages maintained in monoculture or co-cultured with PDCLs from the experiment shown in **(A)** colored by culture condition and annotated using Miller et al. myeloid subtypes. Bar plots show myeloid subtype composition for each PDCL co-culture condition. **(I)** Bar plots showing cell subtype composition for each image in the MILAN study. **(J)** Single cell comparison of *S100A4* RNA expression measured by RNAscope (left) and S100A4 protein intensity measured by COMET staining (right) across annotated cell types in the in-house COMET cohort. **(K)** Amphiregulin secretion in THP-1-macrophage monoculture, PDCL monoculture, and co-culture conditions, measured by nELISA. **(L)** Immunoblots of EGFRvIII expression across GBM PDCLs, including the PDCLs used in the co-culture experiments. **(M)** PDCL viability following treatment with erlotinib plicamycin, or ruxolitinib measured by CellTiter-Glo 2.0. **(N)** Immunoblots and quantification of EGFR, phosphorylated EGFR, STAT3, phosphorylated STAT3, and GSDME expression in CME037 maintained in monoculture with or without erlotinib treatment. **(O)** Immunoblots and quantification of the phosphorylated-to-total STAT3 ratio and GSDME expression following ruxolitinib treatment in the indirect co-culture system. Data are presented as mean ± SEM. Each dot in the bar plots represents an individual replicate. Statistical significance is indicated as *p < 0.05, **p < 0.01, and ****p < 0.0001.

**Supplementary Figure 5.**
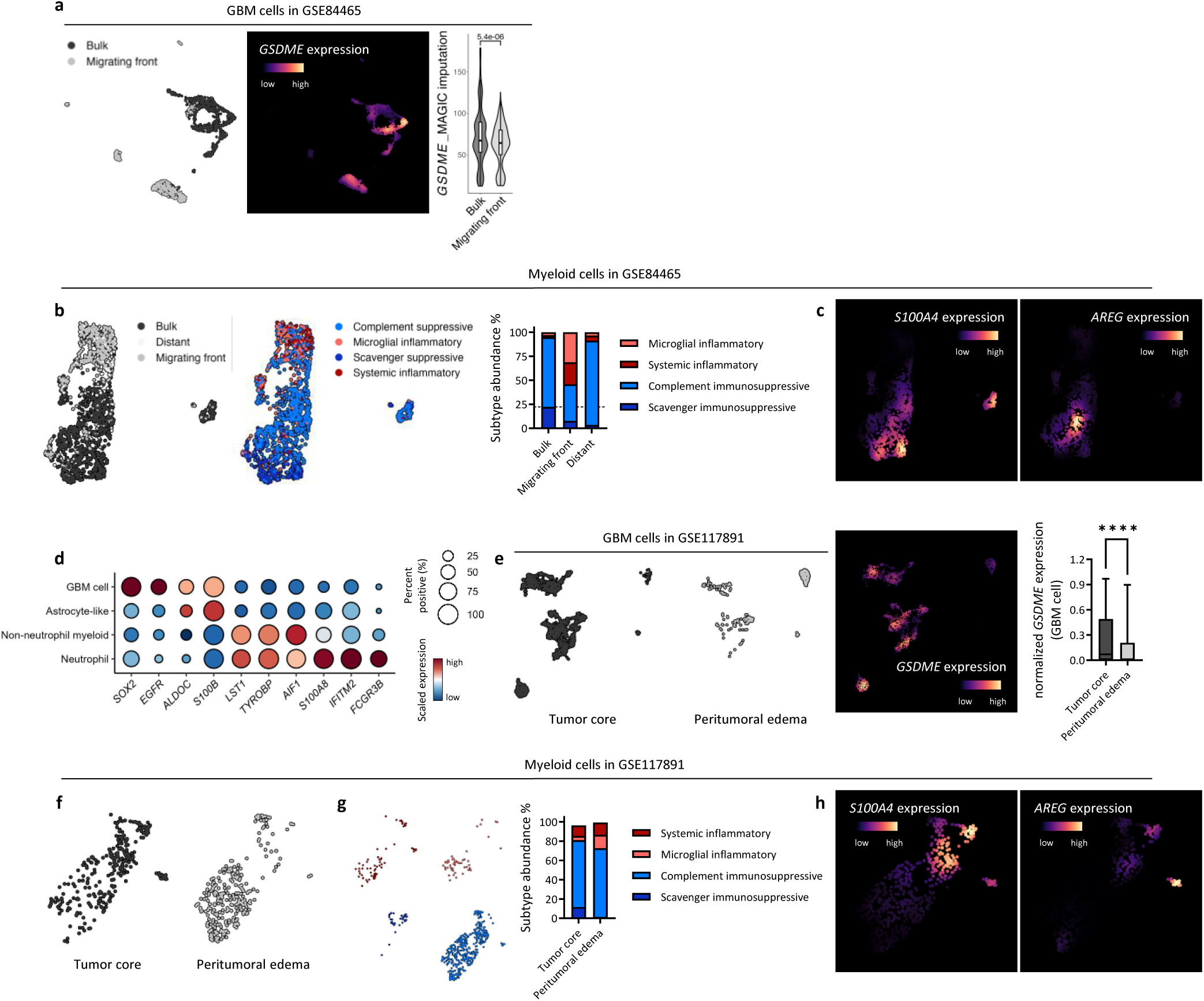
GSDME-expressing GBM cells are enriched in the tumor core. **(A)** UMAP visualization of GBM cells extracted from the scRNA-seq dataset GSE84465. Nebulosa density plot shows GSDME expression, and box plot compares GSDME expression between GBM cells from the tumor core and migrating front. **(B)** UMAP visualization of myeloid cells extracted from the scRNA-seq dataset GSE84465, colored by sampling position (left) and Miller et al. myeloid cellular states (middle). Bar plot shows myeloid subtype composition by sampling position (right). **(C)** Nebulosa density plots showing *S100A4* and *AREG* expression in the myeloid cell UMAP from **(B)**. **(D)** Dot plot showing canonical lineage marker expression across cell types annotated in the GSE117981 scRNA-seq dataset. **(E)** UMAP visualization of GBM cells extracted from the scRNA-seq dataset GSE117981. Feature plot shows GSDME expression, and box plot compares GSDME expression between GBM cells from the tumor core and peritumoral edema. **(F)** UMAP visualization of myeloid cells extracted from the scRNA-seq dataset GSE117981, colored by sampling position. **(G)** UMAP visualization of myeloid cells colored by Miller et al. myeloid cellular states (left). Bar plot shows myeloid subtype composition by sampling position (right). **(H)** Nebulosa density plots showing *S100A4* and *AREG* expression in the myeloid cell UMAP shown in **(F)**. Box plots show the median, interquartile range, and 10^th^-90^th^ percentile whiskers. Statistical significance is indicated as ****p < 0.0001.

**Supplementary Figure 6.**
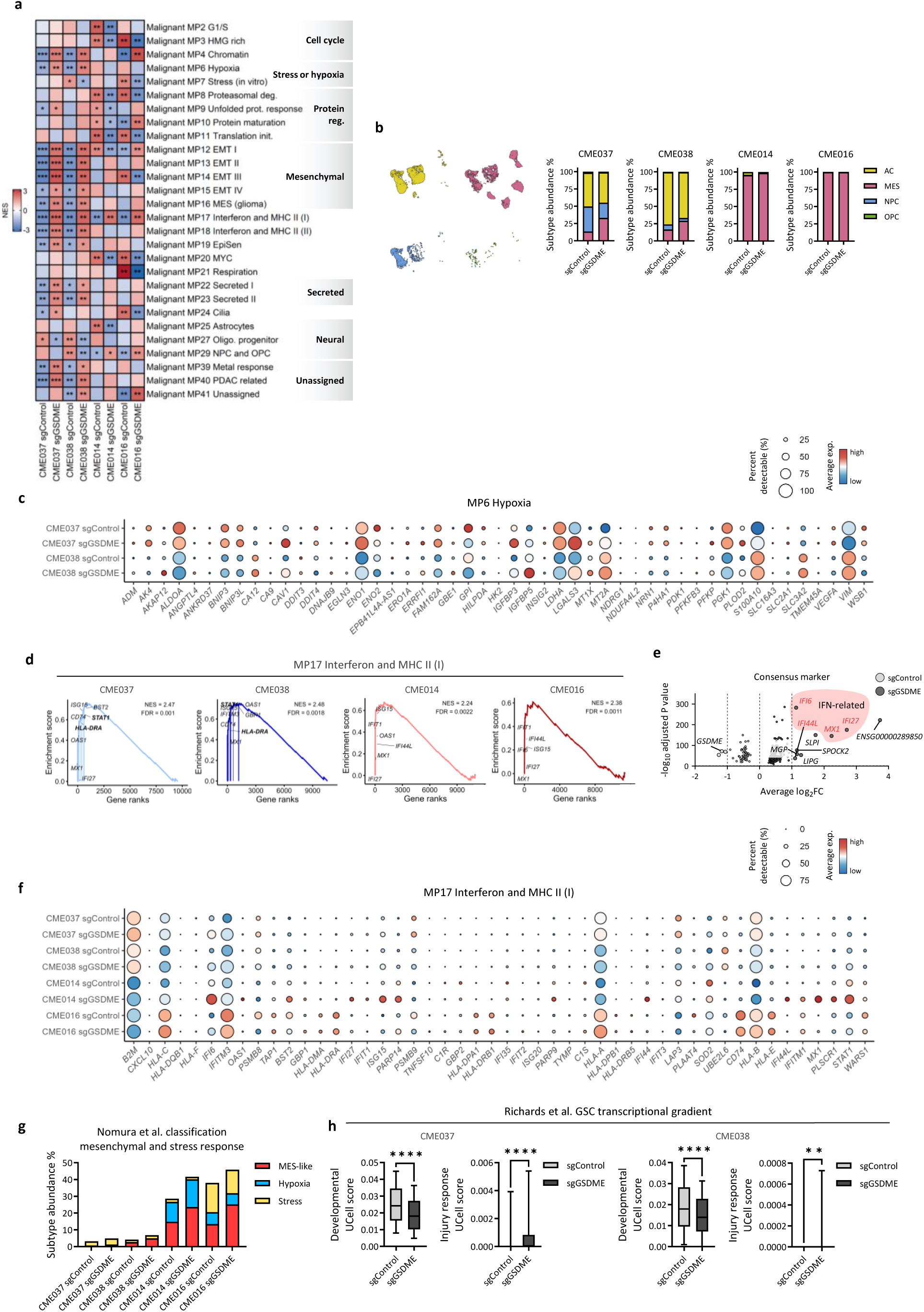
GSDME loss shifts GBM cells toward mesenchymal and hypoxia-associated states. **(A)** Heatmap of GSEA normalized enrichment scores (NESs) for malignant cell metaprograms (MPs) across four GBM PDCLs comparing sgGSDME with sgControl cells. Asterisks within heatmap cells indicate statistically significant enrichment. **(B)** UMAP visualization of scRNA-seq profiles from the GSDME knockout experiment, colored by Neftel et al. GBM cellular states. Barplots show the composition of GBM cellular states in sgControl and sgGSDME cells within each PDCL background. **(C)** Dot plot of expression of the MP6 hypoxia-associated signature. **(D)** GSEA plots of enrichment of MP17, interferon and MHC II (I) in sgGSDME cells across the four PDCLs. **(E)** Volcano plot of consensus upregulated and downregulated genes between sgControl and sgGSDME cells across the four PDCLs, with interferon-related genes highlighted. **(F)** Dot plot of expression of the MP17 interferon and MHC II (I)-associated signature. **(G)** Barplots show the composition of mesenchymal and stress response states in sgControl and sgGSDME cells within each PDCL background. **(H)** Box plots comparing developmental and injury response signature scores between sgControl and sgGSDME cells in CME037 and CME038. Data are presented as mean ± SEM where applicable. Statistical significance is indicated as *p < 0.05, **p < 0.01, and ***p < 0.001.

**Supplementary Figure 7.**
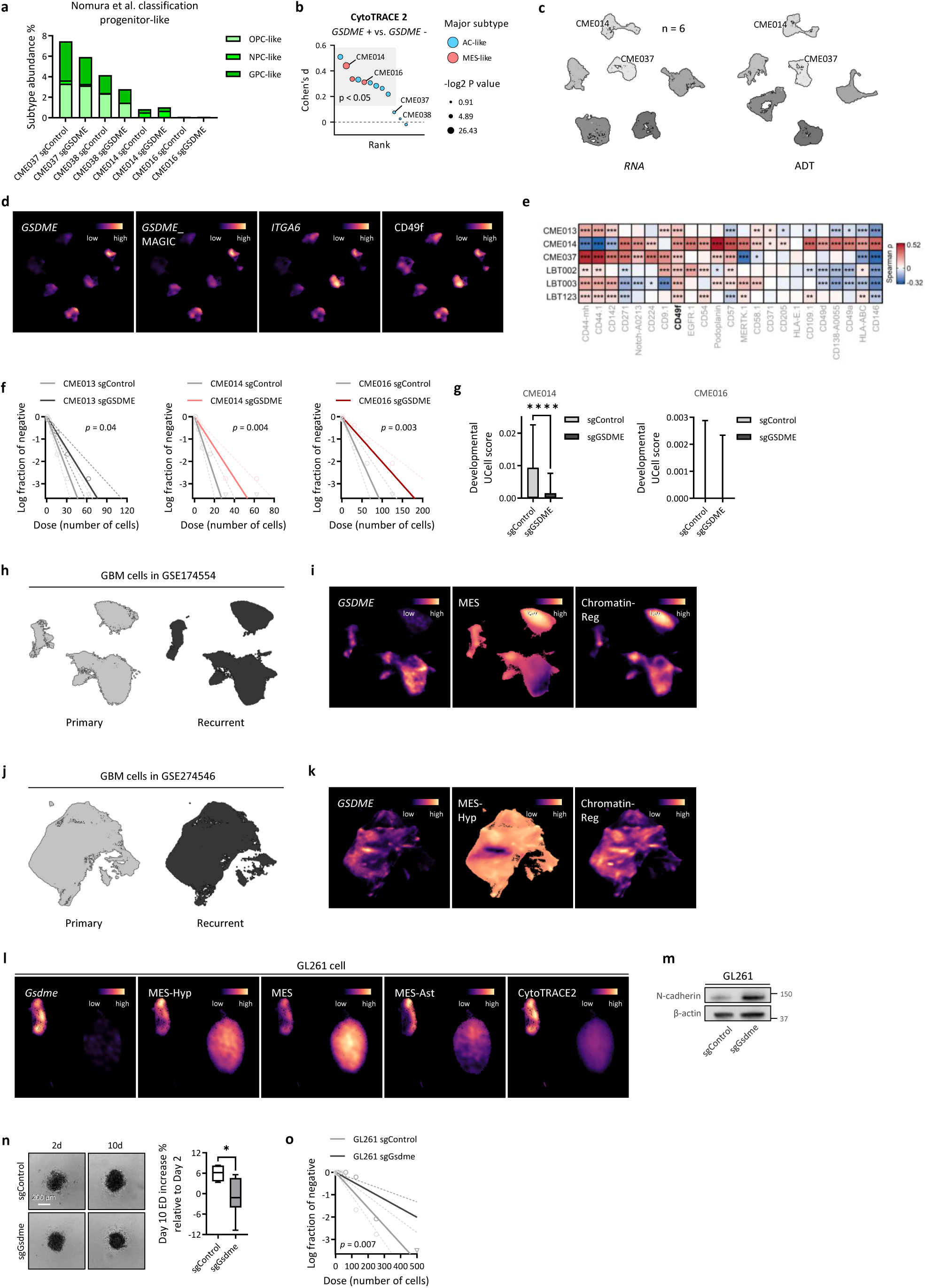
GSDME supports GBM progenitor-like phenotype and is functionally conserved in a murine glioma model. **(A)** Barplots show the composition of progenitor-like states in sgControl and sgGSDME cells within each PDCL background. **(B)** Comparison of CytoTRACE2 scores between *GSDME*^high^ and *GSDME*^low^ GBM cells from the PDCL scRNA-seq dataset. Dot plot shows PDCLs ranked by effect size, with dots colored by the dominant GBM cellular state of each PDCL. **(C)** UMAP visualizations of RNA and ADT profiles from CITE-seq analysis of six GBM PDCLs. **(D)** Nebulosa density plots showing *GSDME* expression, imputed *GSDME* expression, *ITGA6* expression, and surface CD49f protein abundance. **(E)** Heatmap of ADTs upregulated in *GSDME*^hi^ cells in the CITE-seq dataset. Statistical significance (adjusted p value) is indicated as *p < 0.05, **p < 0.01, and ***p < 0.001. **(F)** Box plots comparing developmental signature scores between sgControl and sgGSDME cells in CME014 and CME016. **(G)** Limiting dilution assay comparing sphere formation frequencies between sgControl and sgGSDME cells across multiple PDCLs. **(H)** UMAP visualization of GBM cells extracted from the scRNA-seq dataset GSE174554. **(I)** Nebulosa density plots of cells in **(H)** showing *GSDME* expression, MES score and Chromatin-Reg score. **(J)** UMAP visualization of GBM cells extracted from the scRNA-seq dataset GSE274546. **(K)** Nebulosa density plots of cells in **(J)** showing *GSDME* expression, MES-Hyp score and Chromatin-Reg score. **(L)** Nebulosa density plots of GL261 glioma cell scRNA-seq data showing *Gsdme* expression, MES-Hyp score, MES score, MES-Ast score and CytoTRACE2 score. **(M)** Immunoblots of the mesenchymal marker N-cadherin in sgControl and sgGsdme GL261 cells. **(N)** Gliomasphere formation assay comparing growth of sgControl and sgGsdme GL261 cells. **(O)** Limiting dilution assay comparing sphere formation frequencies between sgControl and sgGsdme GL261 cells. Box plots show the median, interquartile range, and 10^th^-90^th^ percentile whiskers. Statistical significance is indicated as *p < 0.05, and ****p < 0.0001.

**Supplementary Figure 8.**
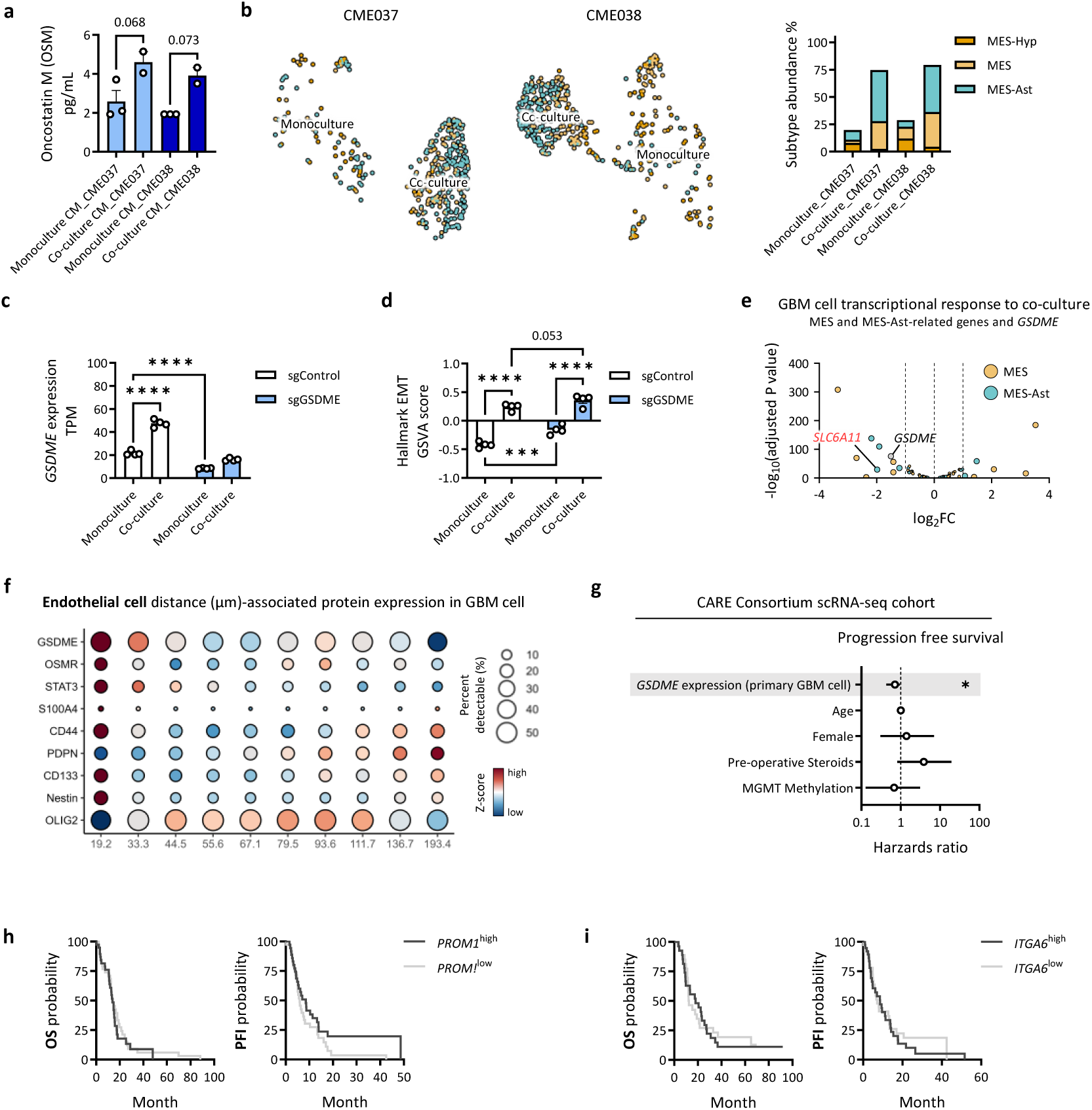
GSDME modulates macrophage-induced mesenchymal transitions in GBM cells. **(A)** Oncostatin M (OSM) secretion in PDCLs monoculture and macrophage co-culture conditions, measured by nELISA. **(B)** UMAP visualization of PDCLs maintained in monoculture or co-cultured with macrophages, colored by Greenwald et al. mesenchymal subtypes (left). Bar plots show the distribution of mesenchymal subtypes across culture conditions for each PDCL (middle), with paired binomial test results shown on the right. **(C)** Bar plot showing *GSDME* expression from bulk RNA-seq of sgControl and sgGSDME CME037 maintained in monoculture or co-cultured with THP-1-macrophages. **(D)** GSVA scores for the Hallmark EMT signature in the bulk RNA-seq conditions shown in **(C)**. **(E)** Volcano plot showing MES and MES-Ast signature genes that overlap with unbiased DEGs between sgControl and sgGSDME GBM cells under macrophage co-culture conditions, after adjustment for monoculture conditions. **(F)** Standardized protein expression per GBM cell across distance deciles relative to endothelial index cells, measured by COMET. **(G)** OS and PFS of GBM patients with *PROM1*^high^ and *PROM1*^low^ tumors in TCGA. **(H)** OS and PFS of GBM patients with *ITGA6*^high^ and *ITGA6*^low^ tumors in TCGA. Data are presented as mean ± SEM where applicable. Each dot in the bar plots represents an individual replicate. Statistical significance is indicated as ***p < 0.001, and ****p < 0.0001.

**Supplementary Figure 9.**
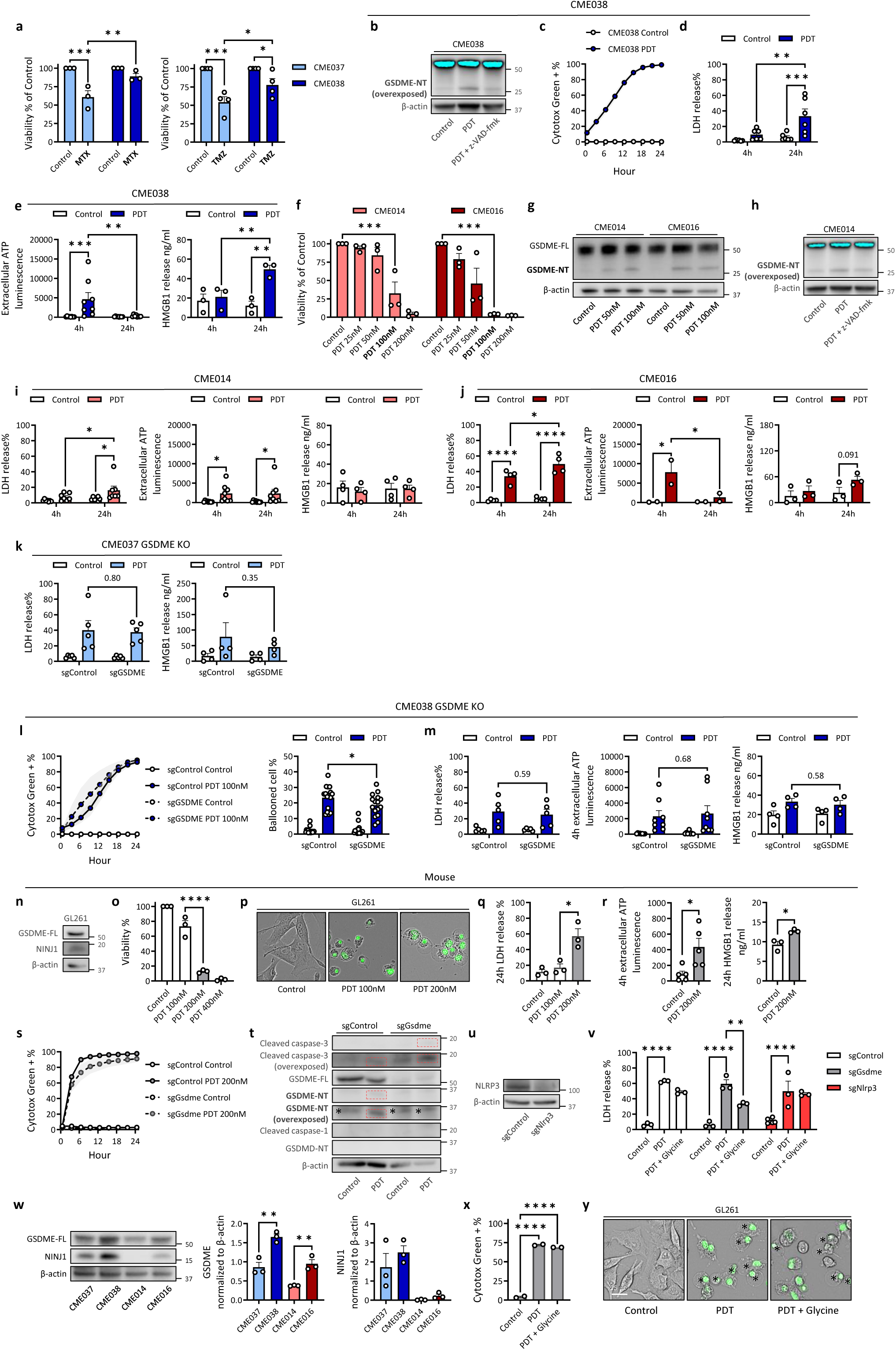
Hyp-PDT induces GSDME-dependent lytic cell death across human and murine glioma models. **(A)** Cell viability of CME037 and CME038 following treatment with MTX or TMZ, measured by CellTiter-Glo 2.0. **(B)** Immunoblot of GSDME cleavage in CME038 after Hyp-PDT in the presence or absence of the pan-caspase inhibitor z-VAD-fmk. **(C)** Time-lapse quantification of Cytotox Green^+^ cells over 24 h in CME038 following Hyp-PDT using the Incucyte live-cell imaging system. **(D)** LDH release at 4 and 24 h in CME038 following Hyp-PDT. **(E)** Extracellular ATP levels and HMGB1 release at 4 and 24 h in CME038 following Hyp-PDT. **(F)** Cell viability of CME014 and CME016 treated with increasing doses of Hyp-PDT, measured by CellTiter-Glo 2.0. **(G)** Immunoblot of full-length GSDME cleavage to the N-terminal fragment in CME014 and CME016 following Hyp-PDT. **(H)** Immunoblot and quantification of GSDME cleavage in CME014 following Hyp-PDT in the presence or absence of the pan-caspase inhibitor z-VAD-fmk. **(I)** LDH release, extracellular ATP levels, and HMGB1 release at 4 and 24 h in CME014 following Hyp-PDT. **(J)** LDH release, extracellular ATP levels, and HMGB1 release at 4 and 24 h in CME016 following Hyp-PDT. **(K)** LDH release (left) and HMGB1 release (right) at 24 h after Hyp-PDT in sgControl and sgGSDME CME037. **(L)** Time-lapse analysis of Cytotox Green positivity in sgControl and sgGSDME CME038 after Hyp-PDT, including the percentage of Cytotox Green^+^ cells over time (left) and quantification of ballooned cell abundance (right). **(M)** Extracellular ATP levels at 4h and LDH and HMGB1 release at 24 h after Hyp-PDT in sgControl and sgGSDME CME038. **(N)** Immunoblot analysis and quantification of GSDME and NINJ1 expression in GL261. **(O)** Cell viability of GL261 treated with increasing doses of Hyp-PDT, measured by CellTiter-Glo 2.0. **(P)** Representative morphology of GL261 cells at 24 h following Hyp-PDT. **(Q)** LDH release after Hyp-PDT in GL261. **(R)** Extracellular ATP levels at 4h and HMGB1 release at 24 h after Hyp-PDT in GL261. **(S)** Time-lapse quantification of Cytotox Green^+^ cells over 24 h in sgControl and sgGsdme of GL261 following Hyp-PDT, acquired using the Incucyte live-cell imaging system. **(T)** Immunoblot of GSDME cleavage and the indicated cell death in sgControl and sgGsdme GL261 following Hyp-PDT. Asterisks indicate nonspecific bands. **(U)** Immunoblots validating CRISPR-Cas9-mediated NLRP3 KO in GL261. **(V)** LDH release at 24 h in the presence or absence of glycine (right) after Hyp-PDT in sgControl, sgGsdme and sgNlrp3 GL261 cells. **(W)** Immunoblot analysis and quantification of GSDME and NINJ1 expression in CME037, CME038, CME014, and CME016. **(X)** Quantification of Cytotox Green^+^ cells at 24 h in GL261 cells under control conditions or following Hyp-PDT in the presence or absence of glycine. **(Y)** Representative morphology of GL261 cells at 24 h following Hyp-PDT in the presence or absence of glycine. Asterisks indicate ballooned cells. Data are shown as mean ± SEM. Each dot in the bar plots represents an individual replicate. Statistical significance is indicated as *p < 0.05, **p < 0.01, ***p < 0.001, ****p < 0.0001.

**Supplementary Figure 10.**
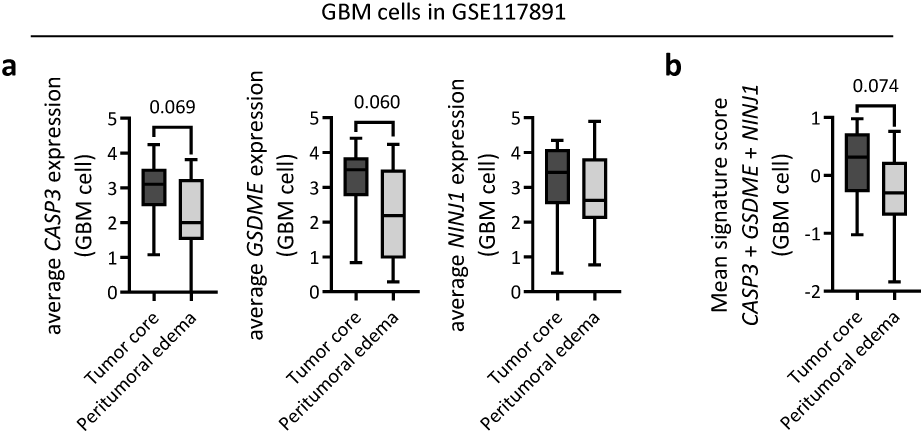
Single cell analysis defines the regional distribution of proinflammatory cell death module in human GBM. **(A)** Box plots of *CASP3*, *GSDME*, and *NINJ1* expression per GBM cell from tumor core versus peritumoral edema samples. Expression values were aggregated across GBM cells within each sample. **(B)** Hyp-PDT signature score in tumor core versus peritumoral edema samples. Box plots show the median, interquartile range and 10^th^-90^th^ percentile whiskers.

